# Editing of *ADA2* Point Mutation in Human Hematopoietic Stem Cells

**DOI:** 10.1101/2025.08.14.670303

**Authors:** Pavel Kopcil, Carolina W. Ervik, Ganna Reint, Katariina Mamia, Monika Szymanska, Shiva Dahal-Koirala, Jacob Conradi, Sigrid Fu Skjelbostad, Oda A. Dønåsen, Xiaojun Jiang, Cecilia Fahlquist-Hagert, Oddrun Kristiansen, Trond M. Michelsen, Espen Melum, Rasmus O. Bak, Anna Komisarczuk, Emma Haapaniemi

**Author notes:** **Corresponding author**:; postal address: Norwegian Centre for Molecular Biosciences and Medicine, Gaustadalléen 21, 0349, Oslo, Norway.

## Abstract

**Background:** The homozygous *ADA2*: c.506G>A (p.Arg169Gln; p.R169Q) variant accounts for majority of Deficiency in Adenosine Deaminase 2 (DADA2). This monogenic disorder may be amenable to *ex vivo* gene therapy by correcting the pathogenic mutation in CD34+ hematopoietic stem and progenitor cells (HSPCs).

**Objective:** To apply CRISPR-Cas9 and homology-directed repair (HDR) as a surrogate strategy to model correction of the pathogenic *ADA2* c.506G>A variant in healthy cord blood HSPCs.

**Methods:** HSPCs were electroporated with optimised CRISPR-Cas9 editing reagents, and editing outcomes, including HDR and on-target deletions, were quantified by ddPCR. Cell functionality was assessed through colony-forming unit (CFU) assays and by xenotransplantation into NOD SCID Gamma (NSG) mice. Two HDR enhancement strategies were tested: (1) genetic inhibitors of p53 and non-homologous end joining (NHEJ) pathways, and (2) pharmacological NHEJ inhibition.

**Results:** Small-molecule NHEJ inhibitors increased HDR efficiency approximately two-fold (from ∼40 % to ∼80 %). Edited HSPCs retained normal CFU capacity and successfully engrafted in NSG mice. However, up to 8 % of edited cells exhibited on-target chromosome loss, though this declined over time. Up to 40 % of T cells and fibroblasts demonstrated similar losses under NHEJ inhibitors treatment. In contrast, genetically encoded inhibitors did not improve HDR.

**Conclusion:** The *ADA2* p. c.506G>A variant can be effectively edited employing surrogate strategy in HSPCs without impairing functionality. Although pharmacological inhibition of NHEJ enhances HDR efficiency, it also increases the risk of on-target chromosome aberrations, highlighting the need for careful consideration of the associated risks and benefits in therapeutic gene editing.

**Key messages:**

1) The *ADA2* p.R169Q variant can be efficiently corrected via HDR, and the edited CD34+ HSPCs retain their engraftment capability in NSG mice.
2) Pharmacological inhibition of NHEJ using small-molecule inhibitors increases HDR efficiency but is associated with significant on-target deletions and chromosomal arm loss, particularly in differentiated cell types, and in a donor-dependent manner.

**Capsule summary:** The *ADA2* p.R169Q variant is a viable target for precision gene editing in hematopoietic stem cells. Although inhibition of NHEJ improves HDR efficiency, it concomitantly increases the risk of large on-target deletions, particularly in differentiated cells.

## Introduction

Deficiency of ADA2 (DADA2) is a monogenic autoinflammatory disorder characterised by various clinical manifestations, including bone marrow failure, systemic vasculitis, ischemic strokes, immunodeficiency, recurrent fevers and skin necrosis^1, 2^. DADA2 is a childhood-onset disease with manifestation before the age of 10. The disease is caused by autosomal recessive mutations in the *ADA*2. The p.R169Q variant (c.506G>A, rs143853578) is particularly enriched in the Finnish population with allelic frequency of 0.1 %^3–5^. Symptomatic medication together with limited bone marrow transplantations prevents application of any life-long curative treatment. CRISPR-Cas9 genome editing offers a promising approach for correcting pathogenic point mutations, including the *ADA2* p.R169Q variant associated with DADA2^6, 7, 8^. In therapeutic applications, patient derived CD34^+^ HSPCs are edited *ex vivo* and subsequently reintroduced via autologous transplantation.

The primary CRISPR-based approaches for point mutation correction include prime editing (PE)^9^, base editing (BE)^10^, and CRISPR-Cas9^7^ used for homology-directed repair (CRISPR/Cas9-HDR). CRISPR/Cas9-HDR can be highly efficient, particularly when combined with pharmacological NHEJ inhibition^11^. However, this method introduces double-strand DNA breaks (DSBs), which carry a risk of genotoxicity^12^. While PE and BE are generally considered to have more favourable safety profiles^13^, prime editing gRNA (pegRNA) design for PE can be cumbersome^14^ and often requires extensive screening of pegRNAs. Importantly, all three methods^15–17^ exhibit locus-dependent variability in both editing efficiency and off-target profile.

In this study, we employed CRISPR/Cas9-HDR to model the correction for *ADA2* p.R169Q variant in cord-blood derived CD34^+^ HSPCs. PE and BE strategies were not evaluated due to their complexity, delivery obstacles and low efficiency in HSPCs^18, 19^ and risk for bystander editing (Fig E1a in the Online Repository)^8, 20^. We demonstrated that *ADA2*-edited cells remain viable and capable of engrafting in immunodeficient mice. However, chromosomal arm deletion (CHAD) at the on-target site were detected in approximately 8 % of edited HSPCs treated with NHEJ inhibitors.

## Methodology

### Ethical statements

All experiments were conducted in accordance with approved ethical permits issued by Norwegian health and research authorities. Healthy donor peripheral blood and mononuclear cells (PBMCs), healthy donor cord blood-derived CD34^+^ hematopoietic stem and progenitor cells (HSPCs), and healthy fibroblasts were obtained under active Research Ethical Committee (REK) applications (ID (2019-2024): 2019/868 and ID (2024-2029): 11481). All animal procedures complied with European Community guidelines and were approved by the Norwegian Food Safety Authority (Mattilsynet) under active FOTS approval (ID 24187; 01.11.2020-31.10.2024). Animal experiments were conducted at the Animal Facility at the Department of Comparative Medicine (KPM), Rikshospitalet, Oslo, Norway.

### Plasmid preparation, cloning and *in vitro* transcription

*Cas9wt, GSE56, Ad5 Orf6&7* and *i53* constructs were cloned into T7-modifed pcDNA-DEST40 backbone to obtain expression vector^21–23^. pCMV-T7-EGFP was a gift from Benjamin Kleinstiver, Harvard University (Addgene plasmid #133962). All plasmids were propagated in *E. coli* and purified using either the MiniPrep (Qiagen) or Maxiprep kit (Thermo Fisher) and sequence-verified via Sanger sequencing (Eurofins).

### Genomic DNA isolation

Genomic DNA was extracted using QIAamp DNA Blood & Tissue Kit (Qiagen). When automated processing was used, the QIAcube HT platform and QIAamp 96 DNA Kit (Qiagen) were employed, following the manufacturer’s instructions.

### Cell culture

All cells were maintained at 37°C, with 21% O_2_ and 5% CO_2_ in a humidified incubator with an open water reservoir. HSPCs were cultured in StemSpan^TM^ SFEM II (STEMCELL Technologies) serum-free medium supplemented with recombinant human TPO, SCF, Flt3-L and IL-6 (Peprotech), in combination with small molecules UM729 and SR-1 (STEMCELL Technologies), and with 1% Pen-Strep (P/S) (Thermo Scientific) and GlutaMax^TM^ (Gibco) (200mM) supplementation ^24, 25^. PBMCs were cultured in RPMI 1640 Medium (Thermo Fisher) supplemented with 10% FBS (Thermo Scientific) and 10% P/S, and recombinant human cytokines IL-2, IL-7, IL-15 (Peprotech), and ImmunoCult^TM^ Human CD3/CD28 T Cell Activator (STEMCELL Technologies)^11^. Fibroblasts were obtained from skin biopsy and cultured in DMEM (Thermo Fisher) supplemented with 10% FBS, 1% P/S. Cells were passaged upon reached 80% confluency^11^.

### Electroporation

Electroporation was performed as previously described^11, 24^. Briefly, Cas9, gRNA and single-stranded oligodeoxynucleotide (ssODN) (IDT) were pre-mixed and combined with cells in electroporation buffer. HSPCs, T cells, and fibroblasts were electroporated using the Lonza 4D-Nucleofector® system with pulse codes DZ-100, EO-115, and CA-137, respectively. For all cells, P3 primary solution was selected. mRNA genetic enhancers were co-electroporated together with editing components.

### NHEJ inhibitors and cells treatment

KU0060648 (TargetMol) and AZD7648 (MedChemExpress) were resuspended in sterile DMSO according to the manufacturer’s instructions when a ready-made version was not available (IDT AltR® HDR Enhancer V2). Cells were placed in the corresponding cell type media without P/S containing NHEJ inhibitor or in media with matched DMSO (Fisher Scientific) concentration. Freshly made media with 1% P/S was added to the cell suspension 24 hrs later.

### Droplet digital PCR

Editing efficiency was quantified using droplet digital PCR (ddPCR) as previously described^11^. Briefly, generated droplets were subjected to amplification using a conventional thermal cycler (Bio-Rad) according to the following protocol: 1) 95 °C – 10 min, 2) 94 °C – 30 sec, 56 °C– 3 min, step repeated 42 times, 3) 98 °C – 10 min, 4) 4 °C – hold. For ddPCR, final reaction volume was 20 µl. Data analysis was performed using QuantaSoft^TM^ or QX Manager software (Bio-Rad). An example of the gating strategy is shown (Fig E2 in the Online Repository).

### Chromosomal arm deletion (CHAD) assay

Chromosomal loss assays were designed to detect long-range deletions and loss of long chromosomal arm distal to the 22q11.1 locus harbouring *ADA2*. A reference probe was designed for *CASC11* gene, located on chromosome 8q24.21. Representative gating examples are shown (Fig E3, 4 in the Online Repository). The percentage of CHAD-positive cells was calculated as follows^12^:

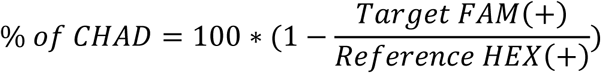

### CFU assays

Semi-solid methylcellulose (STEMCELL Technologies) and liquid (Miltenyi Biotec) CFU assays were performed according to manufacturer’s instructions. The colonies were assessed 14 days after initiation by morphology or flow cytometry using BD® LSR II Flow Cytometer (BD Bioscience). Secondary CFU assay were performed as previously described^26^. All relevant gating plots are shown (Fig E5, 6 and Table 1 in the Online Repository).

### CD34+ staining

For detection of CD34 on the surface of NHEJ inhibitors-treated HSPCs, cells were stained with 7AAD (BioLegend), or Live/Dead Violet (Thermo Fisher) dyes and with CD34-PE (BioLegend) or CD34-APC (BioLegend) antibodies. All cells were blocked with the Human TruStain FcX^TM^ blocker (BioLegend). Samples were analysed on Muse Cell Analyzer (Cytek Biosciences) or on Sony SH800. The data were processed with FlowJo v10.10.0 software (BD Biosciences). Gating strategies are shown (Fig E7, 8 in the Online Repository).

### Humanised mice

8– to 12-weeks-old NOD.Cg-*Prkdc^s^*^cid^ *Il2rg*^tm1Wjl^/SzJ (Charles River) male and female mice were sub-lethally irradiated with two doses of 1.25 Gy, separated by a four-hour interval between each dose. Within 24 hrs post irradiation of the 2^nd^ dose, 200 000 human CD34^+^ HSPCs, resuspended in a total volume of 200 µl sterile PBS, were injected intravenously via the tail vein. Mice were sacrificed 12– or 16-weeks post-irradiation by cervical dislocation. Organs were collected, cells isolated and stained according to the previously established protocols^24,27^. Antibodies and gating plots are shown (Fig E9-14 and Table 2 in the Online Repository). Samples were analysed on BD® LSR II Flow Cytometer (BD Biosciences).

## Results

### Evaluation of established strategies to enhance HDR for ADA2

The local sequence context surrounding the c.506A>G mutation site presents significant challenge for the application of BE due to bystander editing (Fig E1a in the Online Repository). Additionally, *in silico*^14^ predictions for PE yielded suboptimal pegRNAs designs, limiting the feasibility of this approach at the target locus. Consequently, a previously developed and optimised HDR-based genome editing strategy to correct the pathogenic *ADA2* p.R169Q variant^11^ was implemented. The strategy involves an optimised guide RNA (gRNA) and a short single-stranded oligodeoxynucleotide (ssODN) repair template that introduces a wild type SNP (c.506A>G) along with four silent SNPs to prevent re-cutting^28^ by CRISPR-Cas9 and reduce activation of the mismatch repair (MMR) pathway^29^. The HDR protocol was adapted for editing *ADA2* in HSPCs derived from healthy donors (Fig 1a, b). The experimental workflow is depicted in (Fig 1c).

**Figure 1:**
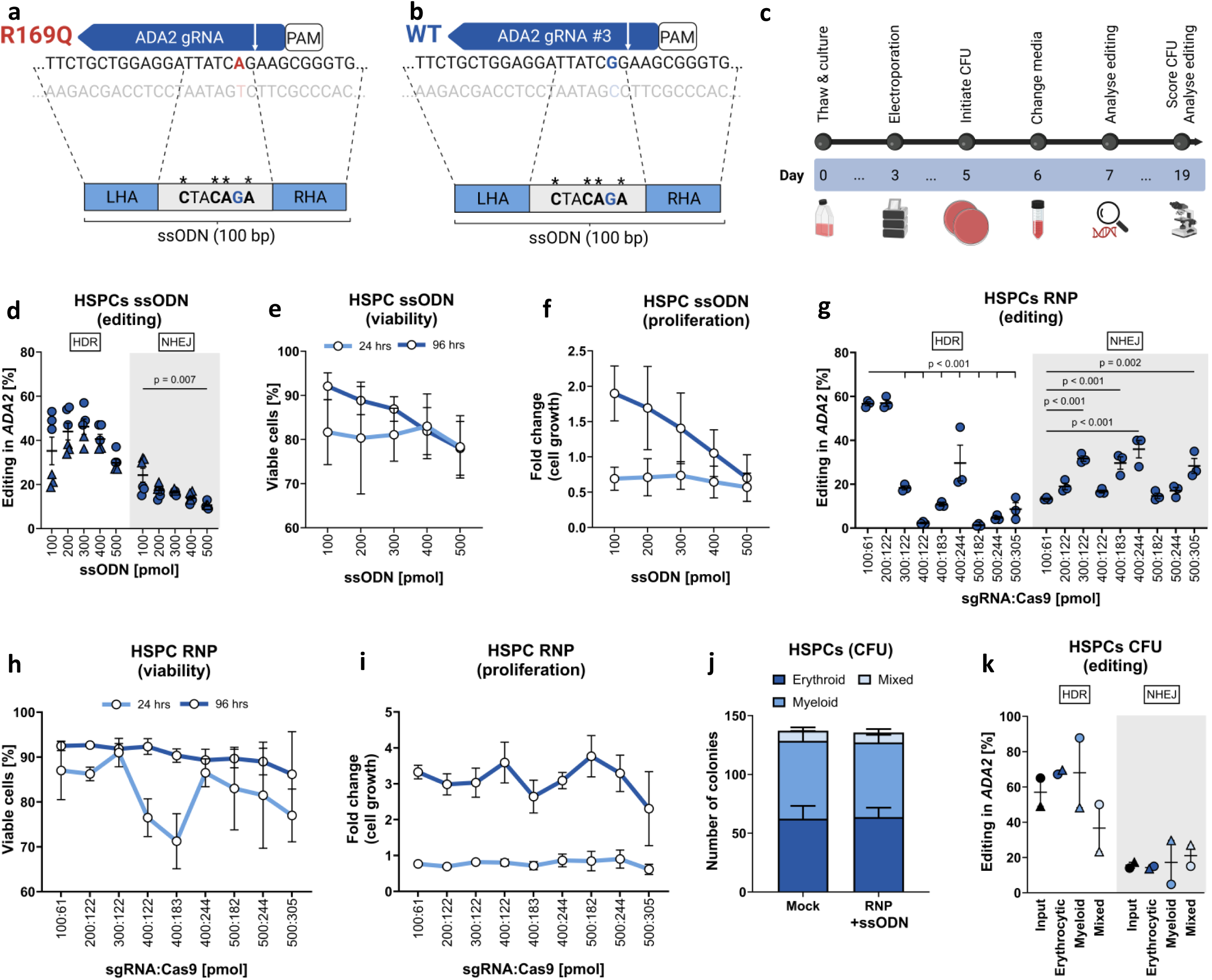
*ADA2* is amenable for CRISPR/Cas9 editing in human CD34^+^ HSPCs. **(a) – (b)** Illustration of the editing strategy for the R169Q mutation or the surrogate strategy targeting the WT *ADA2* gene. **(c)** Schematic timeline of the experimental workflow using cord blood-derived HSPCs. **(d – f)** Editing outcomes, cell viability, and proliferation following ssODN titration. Biological replicates (n=3) in two donors. **(g) – (i)** Editing efficiency, viability and proliferation in response to RNP titration. Biological replicates (n=3) performed using pooled cells from two donors. **(j)** CFU colony classification. Biological replicates (n=2) in three donors. **(k)** Editing efficiency in single-cell-sorted colonies from a semi-solid methylcellulose CFU assay. Biological replicate (n=1) in two donors. Data were analysed either by Two-way ANOVA with Šídák’s (d, e, j), Dunnett’s (g, h, i, k) or Tukey’s (f) multiple comparisons tests. A p-value <0.05 was considered statistically significant.

To determine the optimal editing conditions for the *ADA2* in HSPCs, we conducted a series of titration experiments^11^ using various concentrations of ssODN (Fig 1d-f), and ribonucleoprotein complexes (RNP) (Fig 1g-i). Based on these optimisations, a final condition consisting of 61 pmol Cas9 and 100 pmol of both sgRNA and ssODN was selected.

To ensure the phenotypic identity of HSPCs prior to electroporation, we assessed CD34 surface marker expression after three days of cell culture. Greater than 90 % of CD34^+^ cells was observed (Fig E7 in the Online Repository). To evaluate functional potential of HSPCs post-editing, the colony-forming unit (CFU) assays were performed in semi-solid methylcellulose media. Edited HSPCs formed balanced CFU populations, with editing events detected across all colony types (Fig 1j, k).

Small-molecular inhibitors of NHEJ have previously been shown to enhance HDR^11^. Therefore, we evaluated the effects KU0060648^11^, AZD7648^30^ and IDT HDR Enh. V2^31^ for CRISPR-modified *ADA2* locus (Fig 2a). KU0060648 was used at 0.5µM concentration based on titration experiments (Fig 2b; Fig 15a, b in the Online Repository). For the other two NHEJ inhibitors, 0.5 µM was applied as previously determined concentrations^11, 32^. All inhibitors approximately doubled HDR editing efficiency, achieving ∼80% HDR at the *ADA2* locus (Fig 2c). AZD7648 was the most effective in preserving cell viability, proliferation and CD34 surface expression (Fig 2d-f; Fig E16 in the Online Repository).

**Figure 2:**
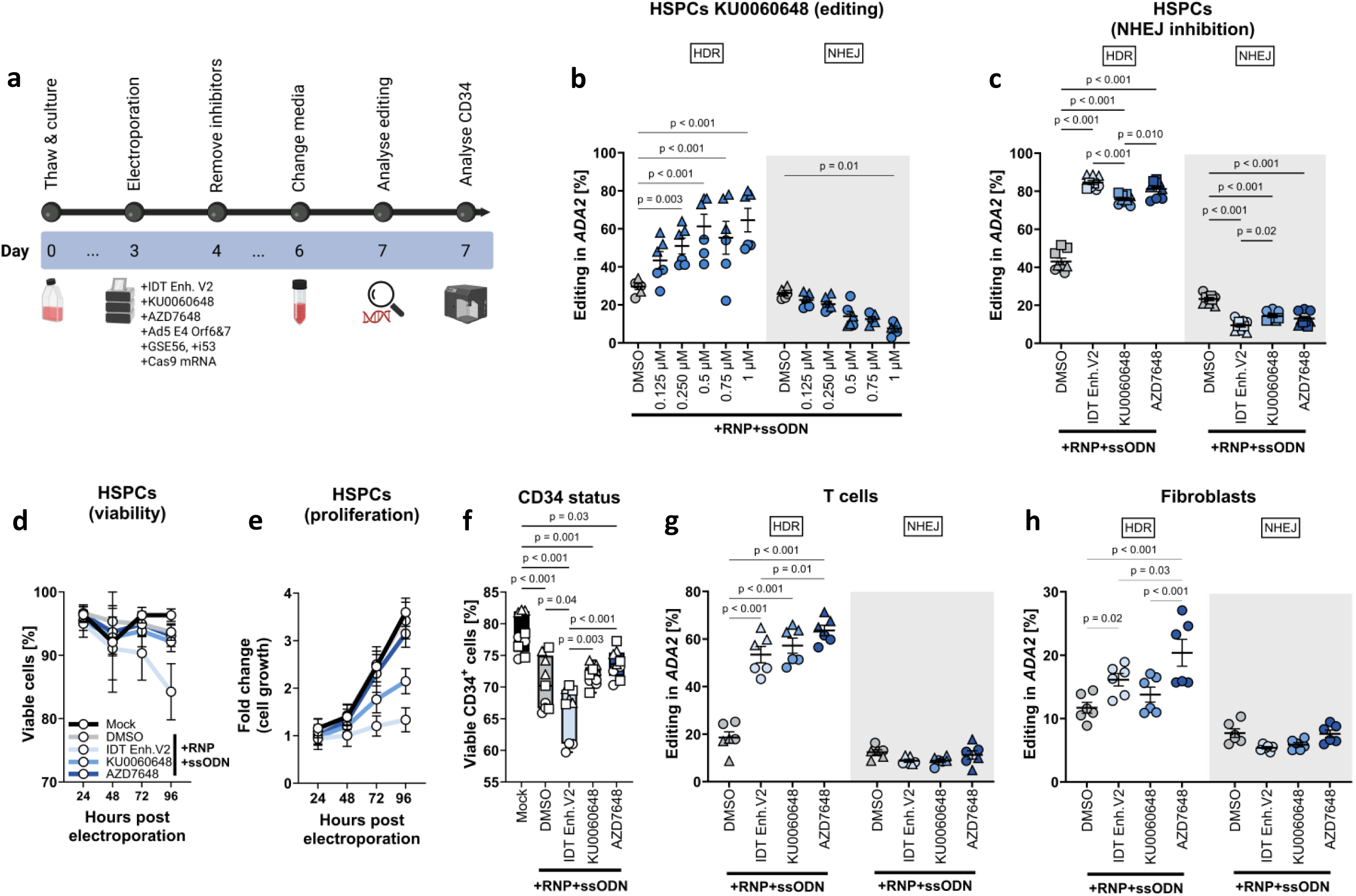
DNA-protein kinase inhibitor AZD7648 elevates HDR outcomes in human primary HSPCs, T cells and fibroblasts. **(a)** Schematic representation of the experimental workflow with small molecule NHEJ inhibitors. **(b)** Editing efficiency of KU0060648 titration in CD34^+^ HSPCs. Biological replicates (n=3), performed in two donors. **(c)** Editing efficiency in CD34^+^ HSPCs treated with NHEJ inhibitors. Biological replicates (n=3) in three donors. **(d) – (e)** viability and proliferation of CD34^+^ HSPCs following treatment with NHEJ inhibitors. Biological replicates (n=3), performed in three donors. **(f)** CD34 surface marker expression in HSPCs treated with NHEJ inhibitors. Biological replicates (n=3) in three donors. **(g)** Editing efficiency in T cells treated with NHEJ inhibitors. Biological replicates (n=3) performed in two donors. **(h)** Editing outcomes in fibroblasts treated with NHEJ inhibitors. Biological replicates (n=3), performed in one donor and repeated twice. Data were analysed by either Two-way ANOVA with Dunnett’s (b) or Tukey’s (c, d, e, f, g, h) multiple comparisons tests. A p-value <0.05 was considered statistically significant.

The inhibitors were also evaluated in healthy donor T cells and fibroblasts. In T cells, all inhibitors yielded 50-60% HDR, a threefold increase over the baseline (Fig 2g; Fig E15c-e in the Online Repository). Fibroblasts exhibited ∼20% HDR with AZD7648, doubling the baseline levels (Fig 2h; Fig 15f in the Online Repository).

Genetically encoded editing enhancers have been reported to increase HDR in HSPCs^22,25^, particularly when the repair template was delivered via adeno-associated viruses^25, 33^. To assess the potential of these enhancers in our ssODN-based editing, we co-electroporated CRISPR reagents with mRNA encoding GSE56 (p53 inhibitor)^22^, Ad5 E4 Orf6&7 (Ad5, inhibitor of inflammatory signaling)^34^ and i53 (NHEJ inhibitor)^21^. However, none of these constructs improved HDR efficiency beyond the levels of EGFP mRNA-treated controls (Fig E17a-c in the Online Repository).

Cas9 fusion proteins have also been reported to enhance HDR ^23, 35, 36^. To evaluate Cas9 fusions in primary immune cells, mRNA or protein-based delivery is required, as plasmids transfection has been reported to trigger an intracellular danger signaling pathway that impairs editing outcomes^37–39^. Therefore, wild type Cas9 mRNA, sgRNA and ssODN were electroporated into HSPCs, T cells, and fibroblasts to assess feasibility of Cas9 mRNA-based editing. Minimal editing was observed in both HSPCs and T cells, whereas fibroblasts obtained editing levels comparable to RNP conditions (Fig E17d-f in the Online Repository). Notably, in the absence of ssODN, Cas9 mRNA triggered robust NHEJ editing (Fig E18 in the Online Repository). Subsequently, we applied several strategies, such as RNAse inhibition and timely delivery of ssODN aiming to improve Cas9 editing efficiency^8, 40, 41^. None of these efforts yielded measurable enhancement in HDR (Fig E19 in the Online Repository).

### NHEJ inhibitors induced chromosomal arm deletions

AZD7648 molecule has been reported to cause on-target CHADs consisting of a chromosomal arm amputations and long deletions in human HSPCs^42^. To evaluate the frequency of CHADs following *ADA2* editing, we designed a ddPCR assay to monitor deletions around the *ADA2* cut site and chromosomal arm amputation distal to chr22q11.1 (Fig 3a).

**Figure 3:**
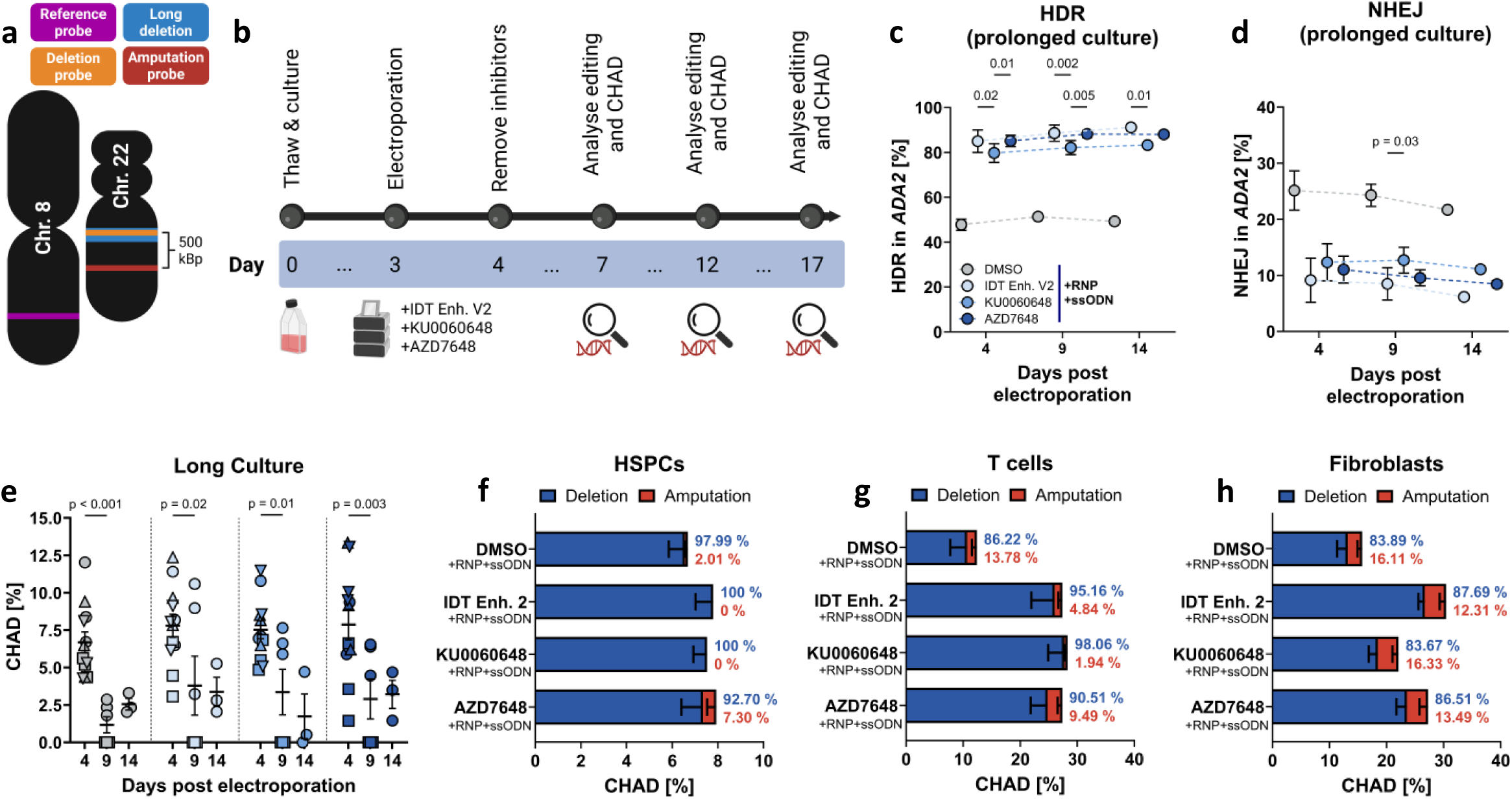
Chromosomal arm deletions in *ADA2*-edited primary cells are prevalent and declines overtime in *in vitro* culture. **(a)** Graphical representation of target and reference locations in the chromosomal arm loss assay. **(b)** Schematic depiction of the experimental design. **(c) – (d)** HDR and NHEJ levels in long-term culture experiment. Biological replicates (n=3), performed in two donors at days five and nine, performed in one donor at day 14. **(e)** Chromosomal arm deletion in long-term culture. Biological replicates (n=3), performed in four donors at baseline, in two donors at day nine and in one donor at day 14. **(f)** Chromosomal arm deletion in T cells. Biological replicates (n=3), performed in two donors. **(g)** Chromosomal arm deletion in fibroblasts. Biological replicates (n=3), performed in one donor repeated twice. Data were analysed by either Two-way ANOVA with Tukey’s multiple comparison test. A p-value <0.05 was considered statistically significant.

We first assessed the presence of CHADs in HSPCs that were cultured for a total of 2 weeks post electroporation (Fig 3b). 80% HDR was observed, that persisted throughout the observation period (Fig 3c, d). Four days post-electroporation, a ∼7% baseline CHAD frequency was detected, which did not increase upon treatment with any NHEJ inhibitors (Fig 3e, f and Fig E20 in the Online Repository).

Next, we evaluated the frequency of CHADs in T cells and fibroblasts. In T cells, CHADs frequencies ranged from 10-20 % across T cell donors (Fig 3g). The NHEJ inhibitors increased the frequency up to 40 %. In fibroblasts, CHADs were approximately 10 % for DMSO and KU0060648, rising to around 30 % with IDT Enh.V2 and AZD7648 (Fig 3h). Our results concludes that CHADs varies by cell type and donor, with terminally differentiated cells being particularly susceptible.

### Comparison of solid and liquid CFU assays to study the effects of NHEJ inhibitors

Genomic rearrangements or DADA2 mutations within *ADA2* can compromise HSPCs biology^43, 44^. The CFU assay serves as an indirect measure of HSPCs differentiation potential. In the solid CFU assay, colony morphology is evaluated by microscopy. In contrast, the liquid CFU assay determines colony phenotype by flow cytometry (Fig E21a, b in the Online Repository).

To compare both CFU methods, live CD34^+^ cells cultured for three days were flow-sorted into 96-well plates containing either solid or liquid media varying in its chemical composition and then cultured for 14 days (Fig E21c in the Online Repository). The solid CFU assay yielded approximately ten times more colonies than liquid assay (Fig E21d in the Online Repository) indicating that single cell sorted HSPCs are not adequately supported by the liquid CFU media. As a result, all subsequent liquid CFU assays were performed using limited dilution.

To investigate the colony-forming capacity of the edited HSPCs, cells were seeded into both solid and liquid plates two days post electroporation (Fig 4a). The colonies were evaluated after 14 days (Fig 4b). To assess the presence of more primitive hematopoietic stem cells in culture, we reseeded the cells from solid CFU colonies into fresh solid CFU media and conducted a second CFU assay (Fig 4c).

**Figure 4:**
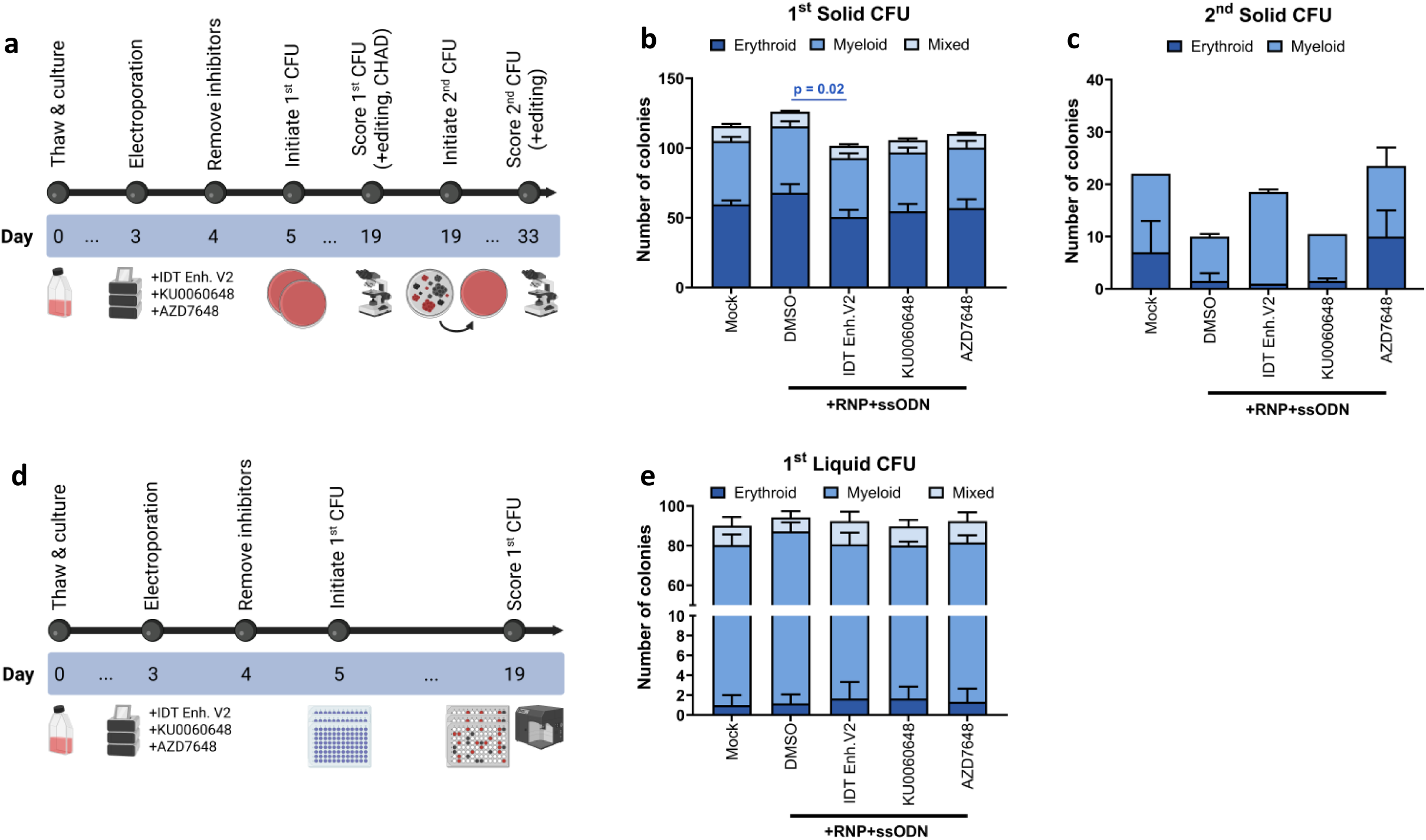
CFU assays shows normal clonal differentiation in *ADA2*-edited HSPCs treated with NHEJ inhibitors **(a)** Experimental design for the solid CFU assay. **(b) – (c)** Colony output in 1^st^ and 2^nd^ solid CFU assay. Biological replicates (n=2) in three donors for 1^st^ CFU, biological replicates (n=2) in one donor for 2^nd^ CFU. **(d)** Experimental design for the liquid CFU assay. **(e)** Colony output in the liquid CFU assay. Biological replicates (n=2) in three donors. Data were analysed by Two-way ANOVA with Tukey’s (b, c, e) and Dunnett’s (b, c, e) multiple comparisons tests. A p-value <0.05 was considered statistically significant.

Comparison of colony types between the solid and liquid CFU assays (Fig 4d), revealed an overrepresentation of myeloid colonies, particularly CFU-Granulocyte (CFU-G; ≥50 % of CD15^+^ cells) in the liquid assay (Fig 4e). Treatment with IDT Enh. V2 reduced the BFU-E output in the solid CFU assay (Fig 4b). In the secondary CFU assay, colony numbers were approximately 10-20% of the initial yield, consistent with previous report^26^. The colony numbers were variable, and the use of inhibitors did not reduce the colony output below control level (Fig 4c).

Genomic DNA was extracted from the colony pools from all experimental conditions to determine HDR editing efficacy and the presence of CHADs (Fig 4a, Fig 5a). HDR rates remained consistent across all treatment groups, ranging from 76 %-85 %, with AZD7648 demonstrating the highest HDR under all experimental conditions (Fig 5b-e). Notably, HDR increases with prolonged culture in AZD7648 treated cells, suggesting effective modification of more primitive HSPCs.

**Figure 5:**
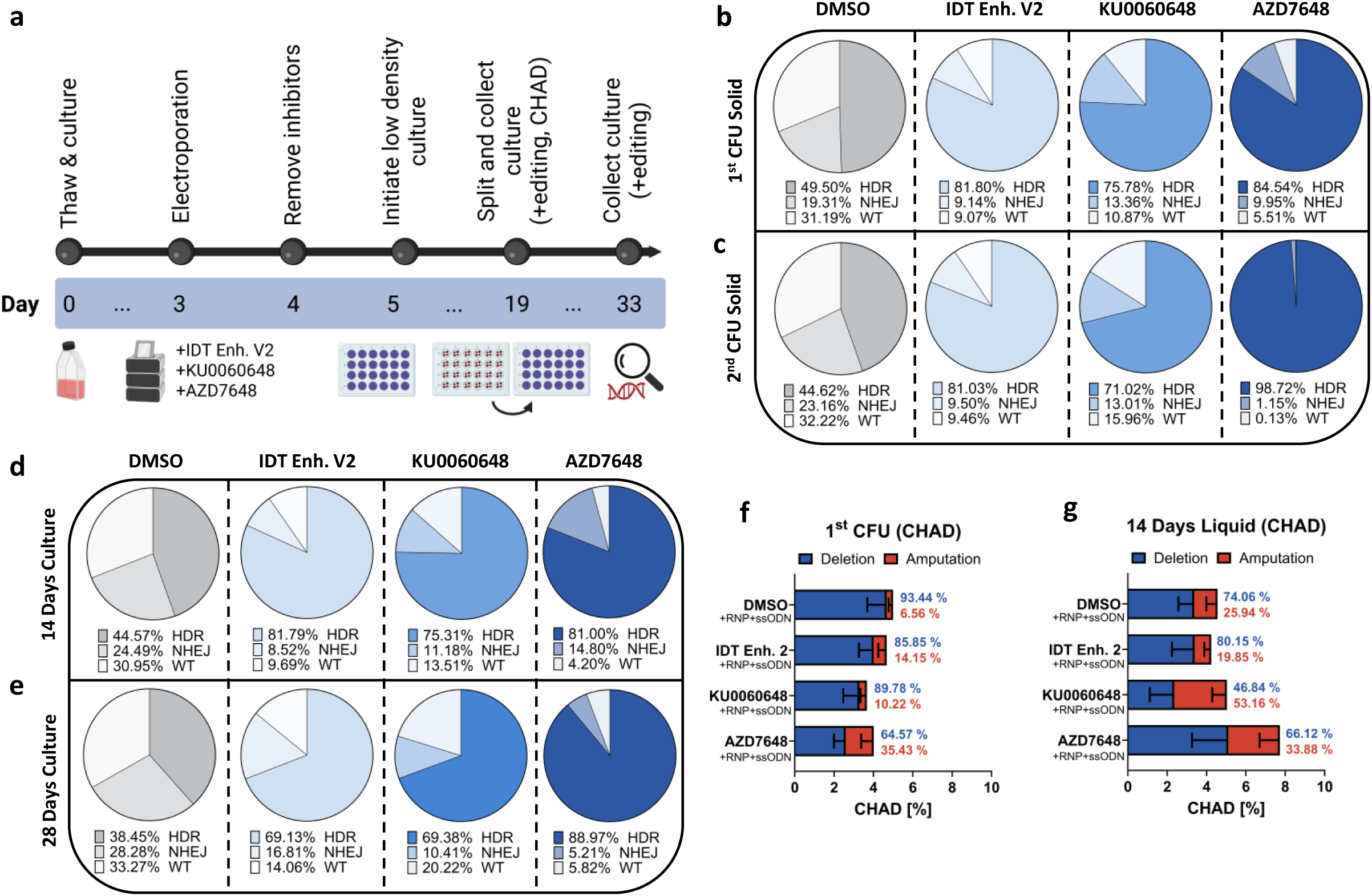
HDR, NHEJ and CHADs are preserved and propagated in both semi-solid and liquid CFU assay. **(a)** Experimental design of the low-density culture in liquid CFU media. **(b) – (c)** Editing outcomes in CFU-differentiated HSPCs from the 1^st^ and 2^nd^ solid CFU assay. Biological replicates (n=2) in four donors for the 1^st^ CFU, in one donor for 2^nd^ solid CFU. **(d) – (e)** Editing outcomes in CFU-differentiated HSPCs after 14 and 28 days in low-density liquid CFU culture. Biological replicates (n=2) in four donors for day 14, biological replicates (n=2) in two donors for days 28. **(f)** CHAD in the 1^st^ solid CFU assay. Biological replicates (n=2) in 4 donors. **(g)** CHAD after 14 days low-density liquid CFU culture. Biological replicates (n=2), performed in four donors. Data were analysed by Two-way ANOVA with Tukey’s multiple comparisons tests. A p-value <0.05 was considered statistically significant.

Finally, we investigated the prevalence of CHADs in cells derived from both CFU assays. Cells from the solid CFU assay demonstrated an average CHAD of ∼5 %, with a trend towards increased amputations in inhibitor groups (Fig 5f). Conversely, the frequency of chromosomal amputations was higher in cells cultured in the liquid CFU media (Fig 5g).

### ADA2-edited HSPCs successfully engrafted in NSG mice

To evaluate whether edited HSPCs retain the capacity to establish a functional bone marrow environment, we transplanted cultured and edited HSPCs into sub-lethally irradiated NSG mice.

Initially, human cell engraftment between cryopreserved unedited, unexpanded (fresh) and cryopreserved HSPCs cultured for four days (expanded) were compared (Fig 6a). Engraftment levels were similar between expanded and fresh HSPCs (Fig 6b). The populations of human CD19^+^, CD33^+^ and CD34^+^ cells (B cells, myeloid cells, and subpopulations of CD34^+^, respectively) were unaffected by the time maintained in culture before transplantation (Fig 6c-e). Subsequently, new cohort of mice were transplanted with *ADA2*-edited HSPCs (Fig 6f). Engraftment levels were similar between edited and unedited cells (Fig 6g), and the retrieved immune cell populations were identical between conditions (Fig 6h-j).

**Figure 6:**
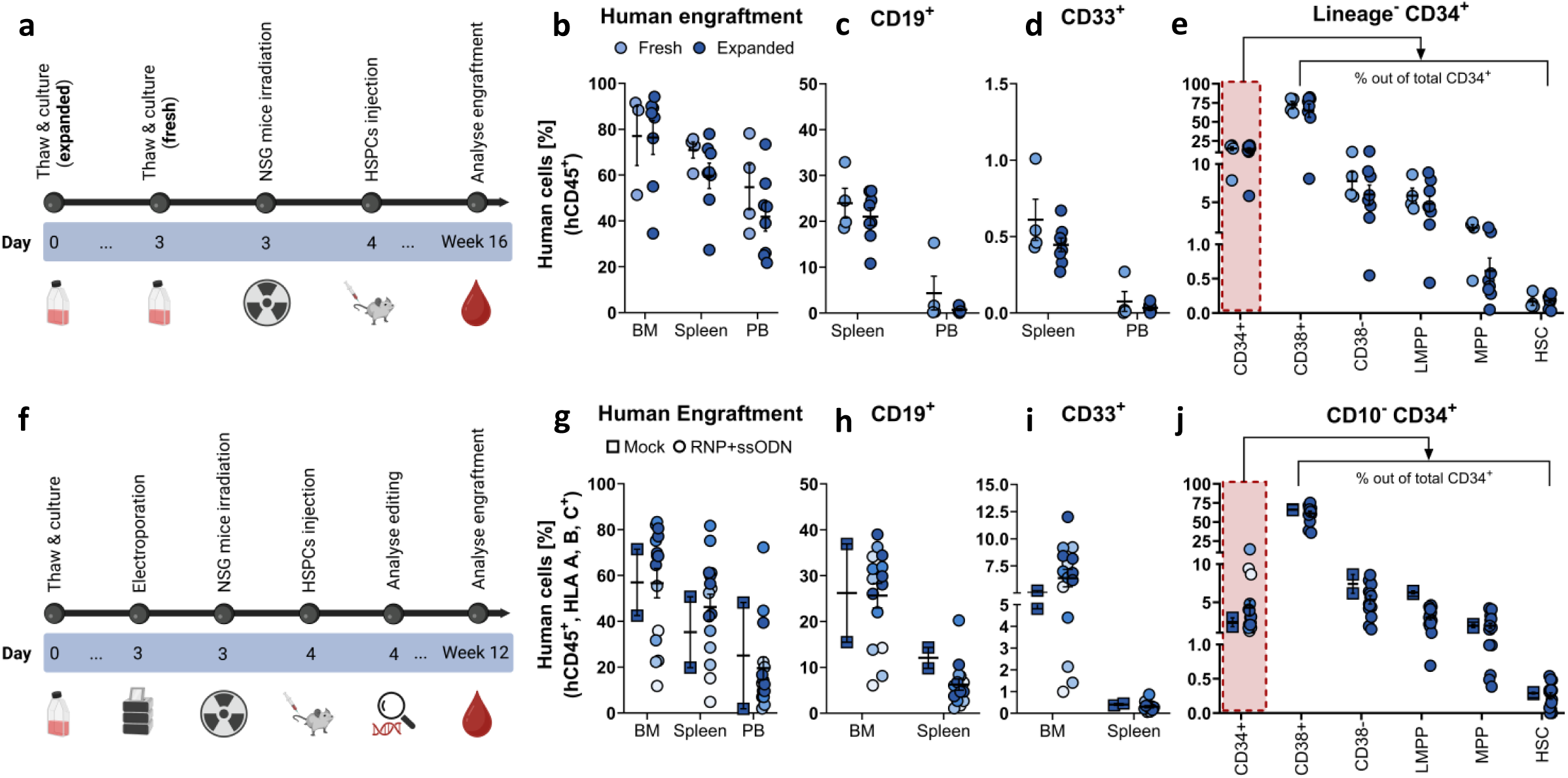
*ADA2* CRISPR/Cas9 modified HSPCs repopulate hematopoietic compartments of NSG mice. **(a)** Schematic timeline of the experimental workflow comparing expanded vs freshly thawn CD34^+^ HSPCs. **(b)** Human engraftment after 16 weeks. Groups: PBS (n=3), Fresh (n=4), and Expanded (n=8) using cells from four donors. **(c) –** (**e)** Percentage of CD19^+^, CD33^+^, and CD34^+^ cell populations. **(f)** Schematic timeline of the experimental workflow using *ADA2* CRISPR/Cas9-edited CD34^+^ HSPCs. **(g)** Human engraftment after 12 weeks. Groups: PBS (n=3), Mock (n=2) and *ADA2* specific RNP+ssODN (n=17), using cells from eight donors, shown in different colours. **(h) –** (**j)** Percentage of CD19^+^, CD33^+^, and CD34^+^ cell populations. **Abbreviations:** BM=bone marrow, PB=peripheral blood; HSC=hematopoietic stem cell, MPP=multi-potent progenitor, LMPP=lympho-myeloid primed progenitor. Both experiments were performed once. Each datapoint represents one mouse. Data were analysed by Two-way ANOVA Šídák’s. A p-value <0.10 was considered statistically significant.

Although overall editing rate remained stable, we observed a trend toward reduced HDR frequency in the bone marrow and spleen samples post-transplantation (Fig 7a). To assess lineages-specific editing, bone marrow-derived human cells from five recipient mice transplanted with cells from the single donor were sorted into CD19^+^, CD33^+^ and CD34^+^ populations. No significant differences were found between the groups (Fig 7b).

**Figure 7:**
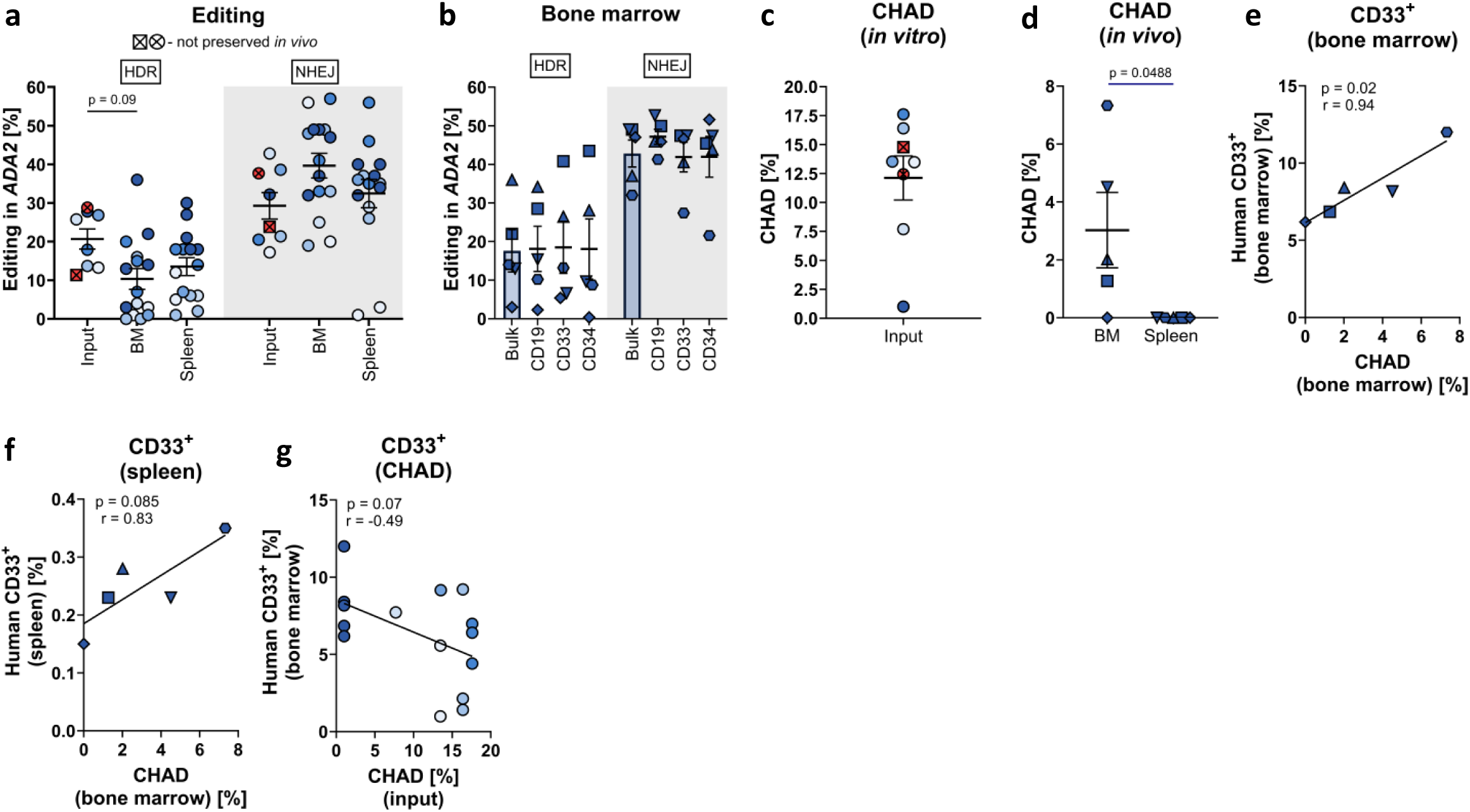
Editing outcomes together with CHAD are preserved in *ADA2* CRISPR/Cas9 modified HSPCs retrieved from humanised NSG mice. **(a)** Editing levels before and after *in vivo* transplantation. For the input, the mean of three replicas from eight donors is shown. Each data point represents one mouse. Symbols with crosses indicates donor cells that were injected but not retrieved from the humanised mice. **(b)** Editing output in sorted cell populations. Each data point represents one mouse (n=5) injected with cells from the same donor. **(c) – (e)** Chromosomal arm deletions before and after *in vivo* period. Performed in duplicates using pooled donor cells for the input and in individual mice (n=5). Mean of the duplicates is shown. **(f)** and **(g)** Correlations between CHAD frequencies in bone marrow (n=5) or input cells (n=15) and the proportion of human CD33^+^ cells retrieved from humanised bone marrow and spleen. Data were analysed by Two-way ANOVA Tukey’s (a, b) multiple comparisons tests, two-tailed unpaired t-test (d) and Pearson multiple variable matrix correlation (e, f, g). A p-value <0.10 was considered statistically significant.

Next, we evaluated the presence of CHADs in the edited, transplanted cells. The pre-transplantation CHAD frequencies were donor-dependent, with an average of ∼12 % across eight donors (Fig 7c). In addition, the CHADs frequency between mice infused with cells from the same donor were compared. While pre-transplanted cells exhibited a baseline CHAD frequency of ∼1 %, the post-transplantation frequency in the bone marrow varied between 0-8%. No CHADs were detected in the spleen, suggesting that bone marrow-specific microenvironmental factors, may support the survival of genomically altered cells (Fig 7d).

In the same cohort, higher CHAD frequencies were discovered in mice with elevated myeloid engraftment, indicating that myeloid progenitors may better tolerate such genomic alterations compared to other lineages (Fig 7e, f). Additionally, we observed a negative correlation between myeloid engraftment and CHAD frequency in the input (pre-transplanted) cells (Fig 7g and Fig E22a, b in the Online Repository).

## Discussion

In this study, we evaluated surrogate CRISPR-Cas9-mediated editing of the *ADA2* p.R169Q founder mutation variant in HSPCs. Using an optimised sgRNA in combination with a chemically modified ssODN and pharmacological inhibition of NHEJ, we achieved on-target HDR efficiencies up to 80 %.

Small molecule inhibitors of the NHEJ pathway have emerged as widely used enhancers of HDR during genomic editing^11, 32, 45^. We compared three previously reported NHEJ inhibitors^31, 46, 47^ and identified that AZD7648 demonstrates the best profile in terms of toxicity and editing enhancement. However, it has been reported that use of AZD7648 during editing can lead to large deletions, particularly in differentiated cells^42, 48, 49^. Consistent with these findings, differentiated cells in our study exhibited deletion rates of up to 40 %, whereas HSPCs demonstrated significantly lower deletion frequency (8%), with minimal increase in response to NHEJ inhibitors. This discrepancy indicates cell-type-specific differences in DNA repair pathway utilisation: differentiated cells tend to rely on error-prone alternative end joining (alt-EJ) when classical non-homologous end joining (c-NHEJ) is inhibited, resulting in large deletions^50, 51^. In contrast, hematopoietic stem cell (HSC) preferentially engages high-fidelity repair pathways and robust DNA damage checkpoints that trigger apoptosis or cell cycle arrest^52, 53^. Thus, we suggest that most HSPCs with lost chromosomal material undergo cell arrest and are gradually selected out from the culture.

In addition to small molecule inhibitors, genetically encoded inhibitors targeting the p53 and NHEJ pathways have been proposed to enhance HDR in HSPCs^22, 25^. We tested mRNA-based delivery of such inhibitors alongside the ssODN but observed no improvement in HDR efficiency in immune cells. We attributed this to the rapid degradation of the exogenous mRNA, potentially due to elevated cytosolic immune sensing and RNase activity in these cells^54–56^. In contrast, fibroblasts tolerated the co-delivery of mRNA and ssODNs^57^. Based on these observations, we conclude that HDR enhancers are most efficient when delivered as proteins or small molecules in ssODN-based systems^32, 58, 59^.

Furthermore, the *ADA2*-edited HSPCs retained the capacity to engraft and reconstitute hematopoiesis in immunodeficient mice, consistent with previous reports involving other loci^17, 60, 61^. Human CD34⁺ HSPCs were recovered from humanised bone marrow post-transplantation confirming successful restoration of hematopoiesis^62^. However, we noticed a modest decline in HDR-edited cells over time^60, 62, 63^, consistent with the hypothesis that long-term repopulating HSCs are less amenable to perform HDR. Future optimisation strategies should aim to increase editing efficiency in this critical subset and to develop culture conditions that support their maintenance and expansion^45, 49, 64, 65^.

We also investigated the presence of CHAD in HSPCs transplanted into NSG mice. Despite a low CHAD frequency at the time of transplantation, the CHAD-positive cells persisted and, in some cases, expanded in the bone marrow. Notably, for *in vivo* experiments we used a chemically unmodified repair template, which is more susceptible to degradation and associated with an increased risk of large deletions^66, 67^. CHAD-positive cells were not detected in the spleen, suggesting that cells harbouring large genomic lesions may depend on the bone marrow microenvironment for survival or are unable to contribute to peripheral hematopoiesis. We found a positive correlation between myeloid engraftment and CHAD frequency, suggesting that myeloid progenitors may be more permissive to chromosomal aberrations than other hematopoietic lineages. Although, clonal expansion was not observed and CHAD did not exceed input levels, more targeted assays—such as barcoding^25^ of HSPCs prior to editing— combined with secondary xenotransplantation experiments^68^, would provide a more precise assessment of the risk of clonal expansion from CHAD-positive cells.

In conclusion, the *ADA2* p.R169Q variant is amenable to CRISPR/Cas9 genome editing in HSPCs. Although NHEJ inhibition substantially improves HDR efficiency, it also increases the risk of on-target chromosomal deletions, particularly in differentiated cells and myeloid progenitors^69–71^. These findings highlight the importance of careful evaluation when applying genome editing strategies to therapeutic contexts. Several inborn errors of immunity, such as DADA2, are associated with an increased risk of myeloid malignancy^72, 73^. Consequently, CRISPR-based therapies may inadvertently accelerate leukemogenesis in a susceptible individual^74–76^. The magnitude of this risk is likely dependent on multiple factors, including the target locus, cell culture conditions, and the individual patient characteristics, underscoring the importance of continued investigation into the safety and context-specific risks of genome editing technologies^48, 49, 77^.

## Declaration of Generative AI and AI-assisted technologies in the writing process

“During the preparation of this work the author(s) used Chat-GPT (OpenAI) in order to improve readability of the presented text. After using Chat-GPT (OpenAI), the author(s) reviewed and edited the content as needed and take(s) full responsibility for the content of the publication.”

## Supporting information

Online Repository Material

HDR to NHEJ Ratios

Individual Datapoints CFU Scoring Editing

Individual Datapoints CHAD

Statistics Main Figures

Statistics Online Repository Figures

Tables Online Repository

## Abbreviations

ADA2: Adenosine Deaminase 2
Ad5: Ad5 E4 Orf6&7
alt-EJ: Alternative End Joining
BE: Base Editing
CFU: Colony Forming Unit
CHAD: Chromosomal Arm Deletion
CRISPR: Clustered Regularly Interspaced Palindromic Repeat
CRISPR/Cas9-HDR: CRISPR coupled HDR
c-NHEJ: Classical NHEJ
DADA2: Deficiency in ADA2
ddPCR: Droplet Digital PCR
DSB: Double Strand Break
HDR: Homology Directed Repair
HSC: Hematopoietic Stem Cell
HSPC: Hematopoietic Stem and Progenitor Cell
NHEJ: Non-Homologous End Joining
NSG: NOD SCID Gamma
IEI: Inborn Errors of Immunity
PE: Prime Editing
pegRNA: Prime Editing Guide RNA
PBMCs: Peripheral Blood and Mononuclear Cells
sgRNA: Single Guide RNA
ssODN: Single Stranded Oligodeoxynucleotide

## Acknowledgements

We are grateful to all donors providing valuable cell material and all hospital personnel assisting during blood collections. We would like to thank Animal Facility (Department of Comparative Medicine, Rikshospitalet, Oslo, Norway), Flow Cytometry Core Facility (Montebello, Radiumhospitalet, Oslo, Norway) and Protein Production Core Facility at Karolinska Institute (Stockholm, Sweden) for using their services and valuable assistance.

## Author contributions

Conceptualization: PK, EH

Methodology: PK, CWE, OK, JC, SFS, OAD, GR, KM, MS, EM, CFH, TMM

Investigation: PK, CWE, JC, SFS, OAD, GR, SDK, JX, MS, CFH

Visualization: PK, CFH

Funding acquisition: EH, TMM

Project administration: PK, MS, AK, EH

Supervision: EH, AK, CFH, ROB

Validation: PK, CWE, JC, SFS, AK

Data curation: PK, CFH, AK, EH

Resources: EH, TMM

Formal analysis: PK, AK, EH

Writing – original draft: PK, AK, EH

Writing – review & editing: PK, CWE, OK, JC, SFS, OAD, GR, KM, SDK, JX, MS, EM, EOB, CFH, TMM, AK, EH

## Data and materials availability

All supplemental and accompanying data are listed as follows:

“Supplementary Material and Methods.pdf”

“Online Repository Material.pdf”

“HDR to NHEJ Ratios.exc”

“Individual Datapoints CFU Scoring and Editing.exc”

“Individual Datapoints CHAD.exc”

“Materials and Methods.exc”

“Statistics_Main Figures.exc”

“Statistics_Online Repository Figures.exc”

“Tables_Online Repository.exc”

Incucyte data storage link: https://osf.io/fkdh6/?view_only=99b31cdffbbe467b917bd60af2f613ae

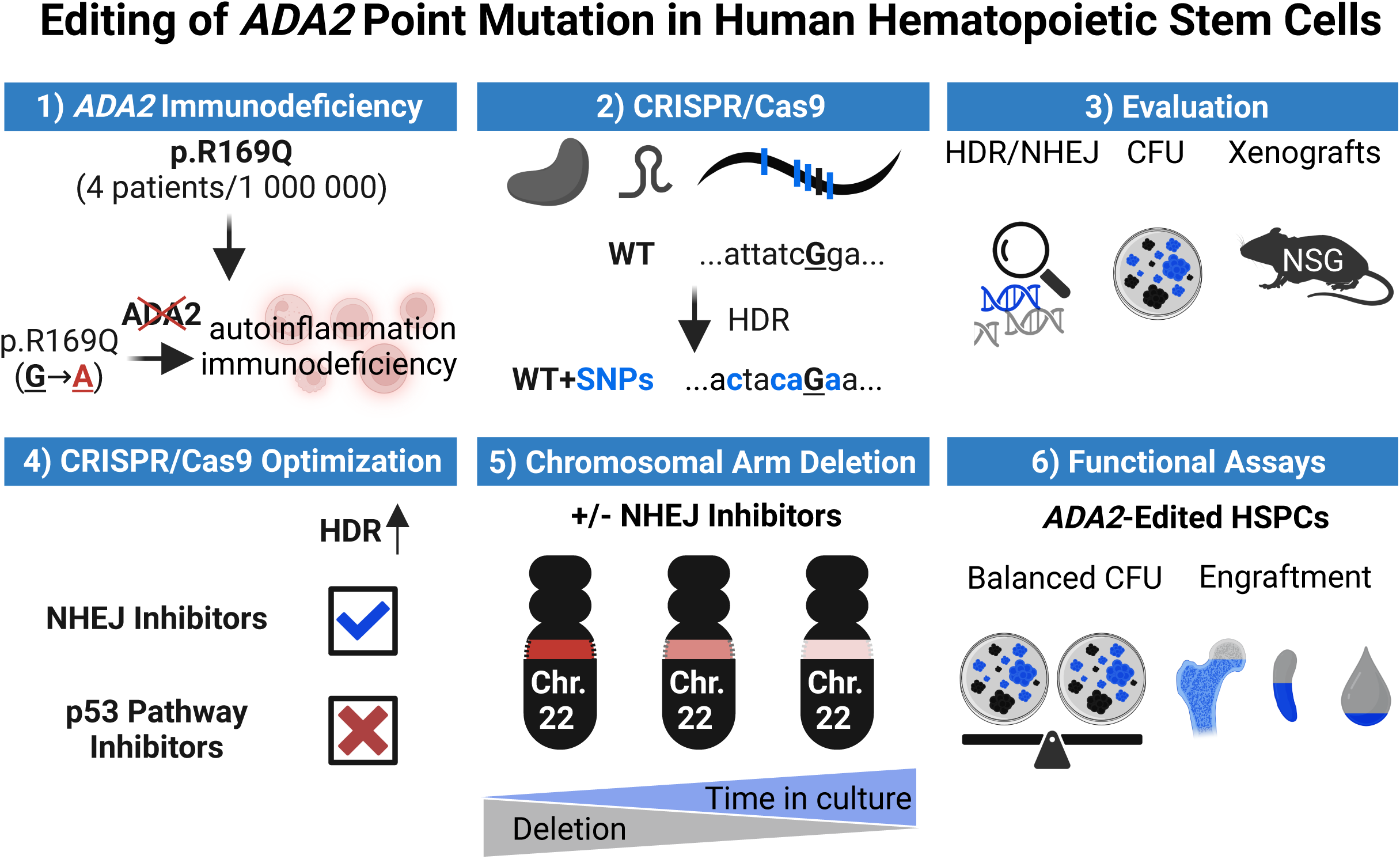

