## Supplementary material for "Editing of *ADA2* Point Mutation in Human Hematopoietic Stem Cells": Online Repository Material

**Online Repository Figure E1: Adenine Base Editing strategy is not feasible for p.R169Q-causing mutation in *ADA2* gene**

Adenine Base Editing (ABE) strategy for correcting pathogenic p.R169Q mutation in *ADA2*. The target nucleotide causing DADA2 is marked in bold red, potential bystander editing sites are depicted in black bold in red circles. The editing window with potential bystander editing sites depends on the variant of adenine base editor.

**Online Repository Figure E2: ddPCR gating examples**

**(a)** Detection of HDR and NHEJ by ddPCR gene editing detection assay. Short ssODN repair template introduces 4 innocent SNPs that matches the HDR detection probe labelled with HEX. A reference probe labelled with FAM is designed to bind to reference sequence that is unaffected by CRISPR/Cas9 (wild type sequence) and is localised outside the ssODN right homology arm region. Total editing (HDR+NHEJ) is detected by utilising drop-off of wild-type probe labelled with HEX binding to unedited wild-type sequence. **(b)** Example of a gating strategy for NHEJ in mock-treated sample. **(c)** Example of a gating strategy for HDR in mock-treated sample. **(d)** Example of a gating strategy for NHEJ in RNP+ssODN-treated sample. **(e)** Example of a gating strategy for HDR in RNP+ssODN-treated sample.

**Online Repository Figure E3: Gating examples of chromosomal arm deletion assay with external reference and loss probe proximal to cut site for three primary cell types, HSPCs, T cells and fibroblasts**

**(a)** FAM labelled probe (in blue) binding to the human *ADA2* sequence downstream of the CRISPR/Cas9 cutting site for detection of chromosomal arm deletions and HEX labelled reference probe (in green) binding in human *CASC11* gene on chromosome 8. Double positive droplets in orange. Gates were set automatically by QX Manager software, ratios between loss and reference values were used to calculate chromosomal arm deletions.

**Online Repository Figure E4: Gating examples of chromosomal arm deletion assay with external reference and loss probe distal to cut site for three primary cell types, HSPCs, T cells and fibroblasts**

**(a)** FAM labelled probe (in blue) binding to the human *BCL2L13* sequence 500 kbp downstream of the CRISPR/Cas9 cutting site for chromosomal arm loss detection and HEX labelled reference probe (in green) binding in the human *CASC11* gene on chromosome 8. Double positive droplets in orange. Gates were set automatically by QX Manager software, ratios between loss and reference values were used to calculate chromosomal arm deletions.

**Online Repository Figure E5: Examples of flow gating for liquid CFU with NHEJ inhibitor-treated HSPCs**

**(a)** Gating strategy and representative FACS plot of a bulk culture to set up gates. Stained and unstained mock samples are shown. **(b)** Representative FACS plots and scoring strategy for each colony type retrieved from liquid CFU assay. A 20% cut-off from the original scoring values was set to control for strict culture duration, ensuring that the number of days in culture neither be extended or shortened.

| **Colony type** | **% of CD15+ cells** | **% of CD14+ cells** | **% of CD235a+ cells** |
|---|---|---|---|
| **CFU-GEMM** | ≥15 | ≥15 | ≥20 |
| **CFU-GM** | ≥30 | ≥30 | |
| **CFU-M** | | ≥50 | |
| **BFU-E** | | | ≥50 |
| **CFU-G** | ≥50 | | |

Table 1: Scoring parameters for liquid CFU assay.

**Online Repository Figure E6: Gating strategy for single cell sorted HSPCs**
**(a)** Gating strategy for single cell sorted HSPCs. Cells were gated for Cells, Single cells, Live cells and $CD34^+$ cells. Sorting was performed into individual 96-well plates by Sony SH800 FACS machine.

**Online Repository Figure E7: FACS gating strategy for cultured $CD34^+$ cells prior electroporation**
**(a)** Gating strategy for unstained sample. **(b)** FACS plots of stained HSPCs. Cells were gated on Cells, Single cells, Live cells and $CD34^+$ cells. Acquisition was performed on SONY SH800 FACS machine. Gating examples are shown for four donors labelled Donor 1 – Donor 4.

**Online Repository Figure E8: Flow charts of NHEJ inhibitors-treated HSPCs for CD34 detection**
**(a)** Schematic representation of gating strategy. **(b)** Gating strategy and representative FACS plots right before electroporation. Mock sample is shown. **(c)** Gating strategy and representative FACS plots 4 days after electroporation. Mock sample is shown. Cells were analysed by Muse and gating was performed in FlowJo (10.10.0).

**Online Repository Figure E9: List of flow cytometry antibodies used for detection of human engraftment in NSG mice.**
**(a)** Overview of flow antibodies used in flow cytometry-related humanised mice experiments. Marker name, clone, catalogue # and antibody types are depicted. **(b)** Schematic tree for gating strategy for two panels utilised for flow cytometry-related humanised mice experiments.

| Flow marker | Clone | Cat. # | Antibody type |
|---|---|---|---|
| PE anti-human CD33 | WM53 | 303404 | Mouse anti-human |
| APC anti-human CD19 | HIB19 | 982406 | Mouse anti-human |
| Alexa Fluor 700 anti-human HLA-A, B, C | W6/32 | 311438 | Mouse anti-human |
| Pacific Blue anti-human CD45 | HI30 | 982306 | Mouse anti-human |
| PE/Cyanine7 anti-mouse CD45.1 | A20 | 110730 | Mouse (A. SW) |
| PE/Cyanine5 anti-mouse TER-119 | TER-119 | 116210 | Rat anti-mouse |
| FITC anti-human Lineage Coctail | RPA-2.10, OKT3, 61D3, CB16, HIB19, TULY56, HIR2 | 22-7778-72 | Mouse anti-human |
| FITC anti-human CD10 | HI10a | 312208 | Mouse anti-human |
| APC anti-human CD34 | 561 | 343608 | Mouse anti-human |
| PE/Cyanine7 anti-human CD38 | HIT2 | 303516 | Mouse anti-human |
| Brilliant Violet 421 anti-human CD90 (Thy-1) | 5E10 | 328122 | Mouse anti-human |
| PE/Cyanine5 anti-human CD45RA | HI100 | 304110 | Mouse anti-human |
| Brilliant Violet 711 anti-mouse CD45.1 | A20 | 110739 | Mouse anti-human |
| PE, anti-human CD45 | HI30 | 304039 | Mouse anti-human |

Table 2: List of antibodies used for analysis of humanised mice.

**Online Repository Figure E10: Gating strategy for FACS panel #2 for humanised bone marrow**
**(a)** Representative gating strategy and FACS plots for bone marrow (FACS panel #2) in humanised mice.

**Online Repository Figure E11: Gating strategy for FACS panel #1 for humanised bone marrow, spleen and peripheral blood**
**(a)** Representative gating strategy and FACS plots for unstained spleen (FACS panel #1) in humanised mice. **(b)** Representative gating strategy and FACS plots for stained spleen (FACS panel #1) in humanised mice. **(c)** Representative gating strategy and FACS plots for unstained peripheral blood (FACS panel #1) in humanised mice. **(d)** Representative gating strategy and FACS plots for stained peripheral blood (FACS panel #1) in humanised mice.

**Online Repository Figure E12: Gating strategy for FACS panel #2 for humanised bone marrow from CRISPR/Cas9 *ADA2*-edited human $CD34^+$ HSPCs**
**(a)** Gating strategy and representative FACS plots for unstained bone marrow in humanised mice for FACS panel #2. **(b)** Gating strategy and representative FACS plots for stained bone marrow in humanised mice for FACS panel #2.

**Online Repository Figure E13: Gating strategy for FACS panel #1 for humanised bone marrow, spleen and peripheral blood from CRISPR/Cas9 *ADA2*-edited human $CD34^+$ HSPCs**
**(a)** Gating strategy and representative FACS plots for stained bone marrow in humanised mice for FACS panel #1. **(b)** Gating strategy and representative FACS plots for stained spleen in humanised mice for FACS panel #1. **(c)** Gating strategy and representative FACS plots for stained peripheral blood in humanised mice for FACS panel #1. **(d)** Gating strategy and representative FACS plots for unstained spleen in humanised mice for FACS panel #1.

**Online Repository Figure E14: Sorted human cells retrieved from humanised bone marrow of NSG mice**
**(a)** Gating strategy and representative FACS plots for sorted humanised bone marrow cells. Retrieved cells were sorted into live $CD19^+$, $CD33^+$, $CD34^+$ and $CD3^+$ populations.

**Online Repository Figure E15: DNA-PKcs inhibitor AZD7648 preserves cell qualities in T cells**
**(a)** and **(b)** viability and proliferation of KU0060648 titration in $CD34^+$ HSPCs, biological replicates (n=3), performed in two donors. **(c)** Schematic representation of experimental workflow in T cells for genetic inhibitors of the p53 pathway and NHEJ inhibitors. **(d)—(e)** Viability and proliferation of T cells treated with NHEJ inhibitors, n=3 performed in two donors. **(f)** Schematic representation of experimental workflow in T cells for genetic inhibitors of the p53 pathway and NHEJ inhibitors. Data were analysed by Two-way ANOVA with Dunnett's (a, b) or Tukey's (d, e) multiple comparisons tests. P-value <0.05 was considered significant.

**Online Repository Figure E16: Proliferation of HSPCs monitored with Incucyte**
**(a)** Phase objects count values (nine foci per 96-well) every 24 hrs, **(b)** Phase objects count values (nine foci per 96-well) every 12 hrs. Performed in one donor. **(c)** Phase objects count values (nine foci per 96-well) every six hrs. Biological replicates (n=3), performed in one donor.

**Online Repository Figure 17: Alternative strategies for HDR enhancement are ineffective when delivered as mRNA**

**(a) – (c)** Editing outcomes of inhibition of the p53 pathway in HSPCs, T cells and fibroblasts, biological replicates (n=3), performed in two donors for HSPCs, T cells and in one donor for fibroblasts repeated three times. **(d) – (f)** Editing efficiency in HSPCs, T cells and fibroblasts for Cas9 protein or Cas9 mRNA edited with or without AZD7648, biological replicates (n=3), performed in two donors for HSPCs, T cells and in one donor for fibroblasts. Data were analysed either by Two-way ANOVA with Tukey's multiple comparisons test. P-value <0.05 was considered significant.

**Online Repository E18: mRNA-based strategies for HDR improvement and Cas9 mRNA-based approach compromise editing when short ssODN ss employed as a repair template**

**(a)** Co-electroporation of corresponding and non-corresponding chemically unmodified ssODN towards *ADA2* site and **(b)** effect of different capping structure in Cas9 mRNA, biological replicates (n=3), performed in one donor. **(c)** Editing and **(d)** viability in T cells with chemically modified (3' 2xPT) ssODN, biological replicates (n=3), performed in one donor. **(e)**

**Online Repository Figure E19: Two-Step electroporation and RNAse inhibition does not improve editing with Cas9 mRNA**

**(a)** Schematic of experimental design for Two-Step Electroporation or RNAse Inhibition in T cells. **(b)** Two-Step Electroporation and **(c)** RNAse Inhibition editing outcomes in T cells, biological replicates (n=3), performed in three donors (Two-Step Electroporation) or in two donors (RNAse Inhibition). **(d) – (g)** Viability and proliferation for Two-Step Electroporation and RNAse Inhibition in T cells, n=3 in three donors for Two-Step electroporation, in two donors for RNAse Inhibition. Data were analysed by Two-way ANOVA with Tukey's multiple comparisons tests. P-value <0.05 was considered significant.

**Online Repository Figure E20: Pearson correlation coefficient (r) for NHEJ inhibitor treated HSPCs in baseline and CFU assays.**

**(a)** DMSO-treated HSPCs for solid and liquid CFU media. **(b)** IDT HDR Enhancer V2-treated HSPCs for solid and liquid CFU media. **(c)** KU0060648-treated HSPCs for solid and liquid CFU media. **(d)** AZD7648-treated HSPCs for Solid and Liquid CFU media.

**Online Repository Figure E21: Cell morphology and phenotypic classification of CFU colonies derived from solid and liquid CFU assays.**

**(a)** Representative photographs of individual colonies retrieved from semi-solid methylcellulose CFU assay. Photographs were taken either by microscope camera or by Incucyte imaging system. **(b)** Representative illustration of individual colonies obtained from liquid CFU assay by photography and flow cytometry plots. Photographs were taken by Incucyte imaging system. **(c)** Experimental schematic of single cell sorted CFU assays. **(d)** Retrieval efficacy after single cell sorted HSPCs either into solid or liquid CFU media, performed in 96-well plate (n=2) in two donors. Data were analysed by unpaired t-test, P-value <0.05 was considered significant.

**Online Repository Figure E22: Pearson correlation heatmap for engrafted *ADA2*-edited HSPCs**

**(a)** Pearson correlation (r) coefficient represented in multiple variable matrix heatmap for data relationship between 15 mice injected with six HSPCs donors. **(b)** Pearson correlation (r) coefficient represented in multiple variable matrix heatmap for data relationship between five mice injected with one HSPCs donor. Statistical significance with p-values is shown in Excel file with all statistical values.

**Online Repository Figure E23: Gel pictures of linearised plasmids and *in vitro* transcribed mRNA**

**(a)** and **(b)** plasmids of genetic inhibitors of the p53 pathway and mRNA products of genetic inhibitors of p53. **(c)** and **(d)** plasmids of the Cas9 wild-type and the mRNA product of Cas9.

**Online Repository Figure E24: ddPCR gene expression gating examples for *p21*, *IFNB1* and *OAS2* assays.**

**(a)** Gating example for *p21*, reference (HEX, *GAPDH*) gene in green, *p21* (FAM) gene in blue. Double-positive droplets are shown in orange. **(b)** Gating example for *IFNB1*, reference (HEX, *GAPDH*) gene in green, *IFNB1* (FAM) gene in blue. Double-positive droplets are shown in orange. **(c)** Gating example for *OAS2*, reference (HEX, *GAPDH*) gene in green, *OAS2* (FAM) gene in blue. Double-positive droplets are shown in orange.

a

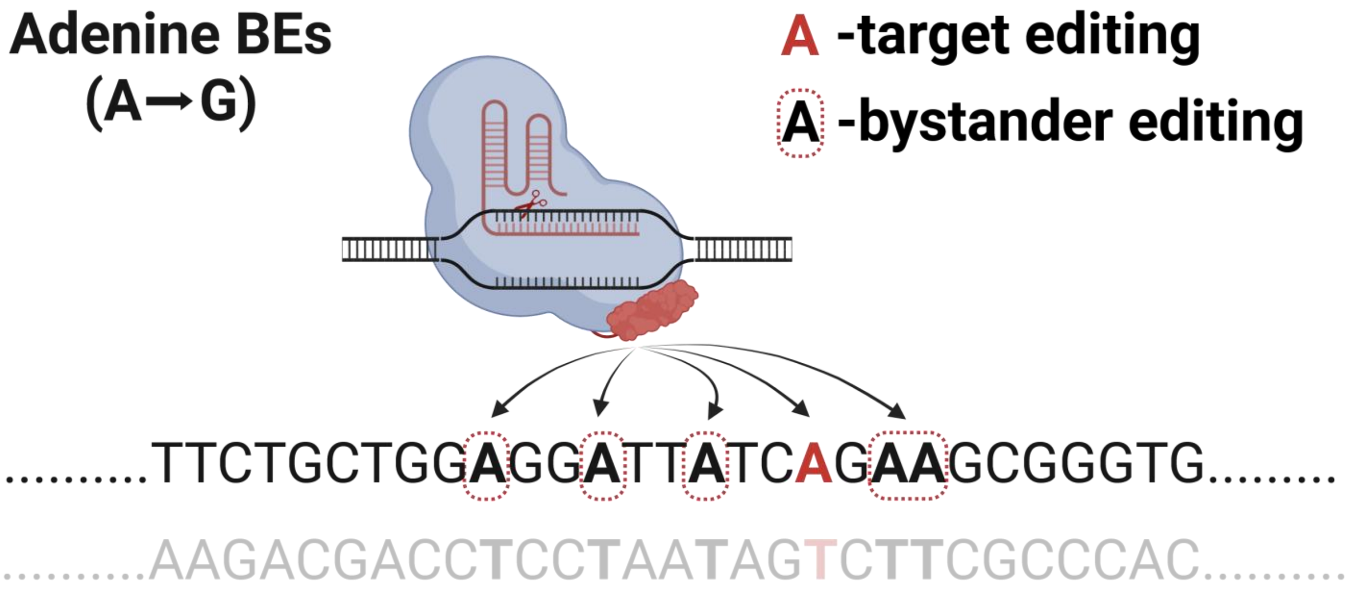
Adenine BEs
(A→G)
A -target editing
A -bystander editing
..........TTCTGCTGGAGGATTATCAGAAGCGGGTG.........
..........AAGACGACCTCCTAATAGTCTTCGCCCAC..........

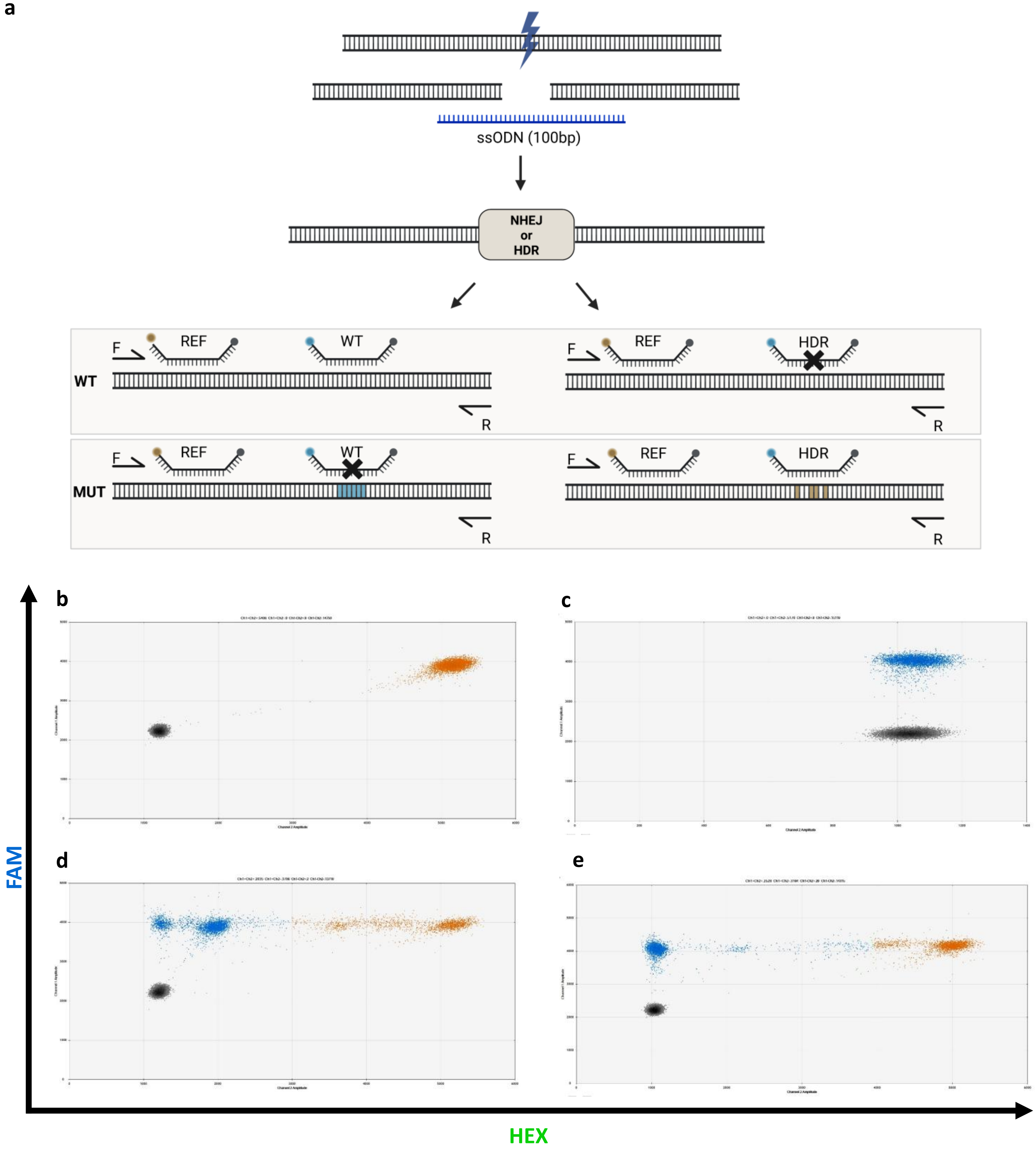
a
ssODN (100bp)
NHEJ
or
HDR
F
REF
WT
R
HDR
MUT
b
c
d
e
FAM
HEX

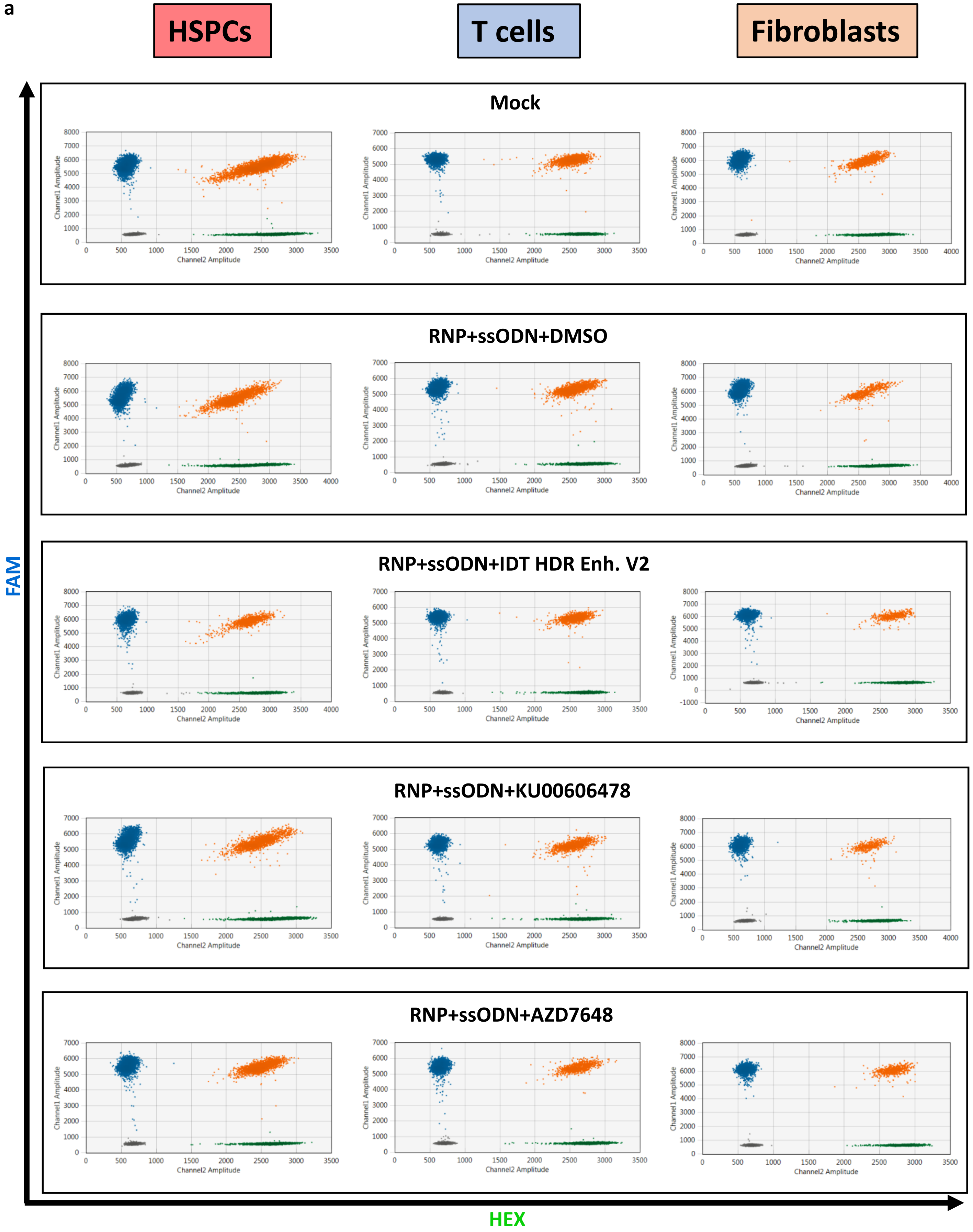
a
HSPCs
T cells
Fibroblasts
Mock
RNP+ssODN+DMSO
RNP+ssODN+IDT HDR Enh. V2
RNP+ssODN+KU00606478
RNP+ssODN+AZD7648
Channel1 Amplitude
Channel2 Amplitude
FAM
HEX

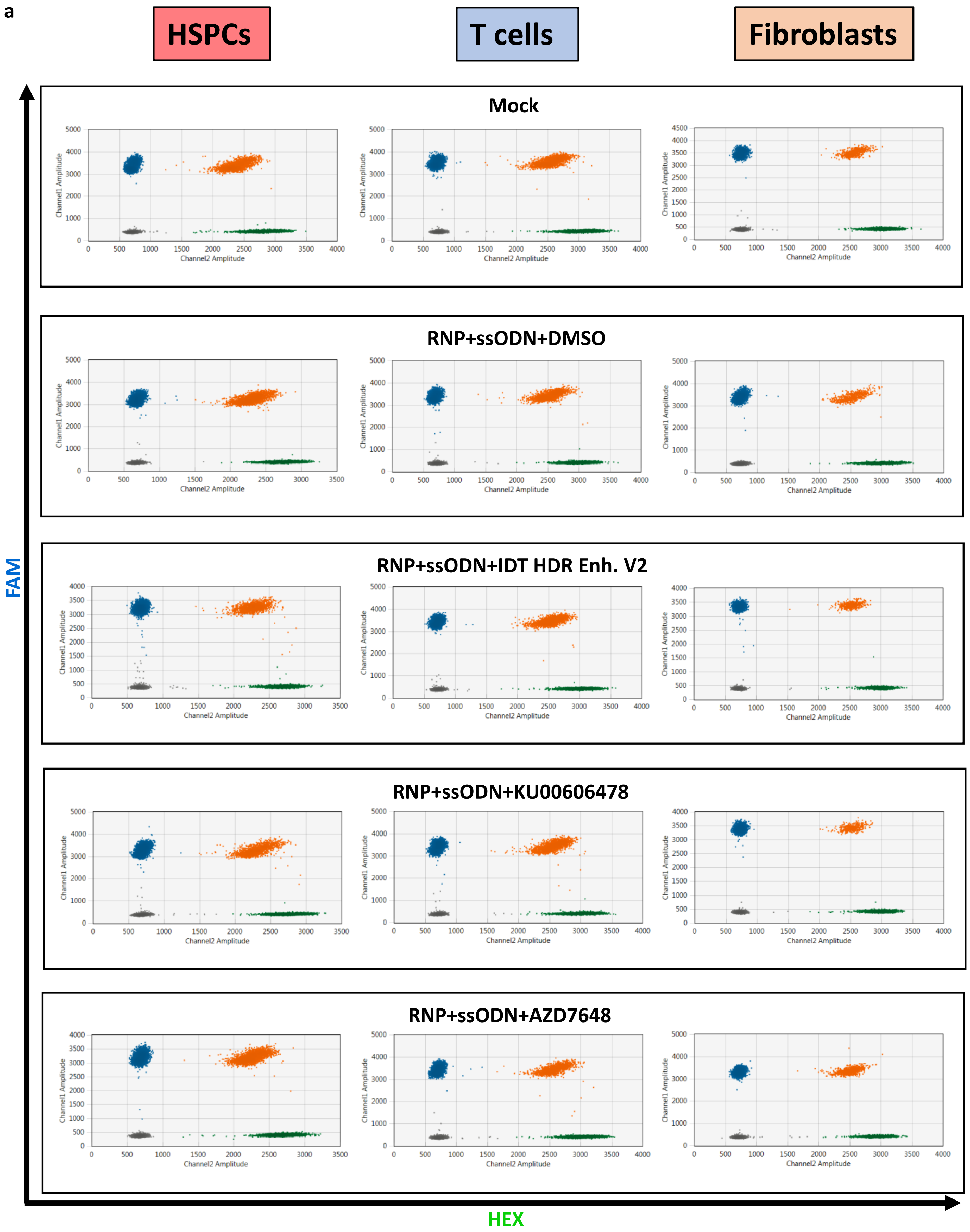
a
HSPCs
T cells
Fibroblasts
Mock
RNP+ssODN+DMSO
RNP+ssODN+IDT HDR Enh. V2
RNP+ssODN+KU00606478
RNP+ssODN+AZD7648
Channel1 Amplitude
Channel2 Amplitude
FAM
HEX

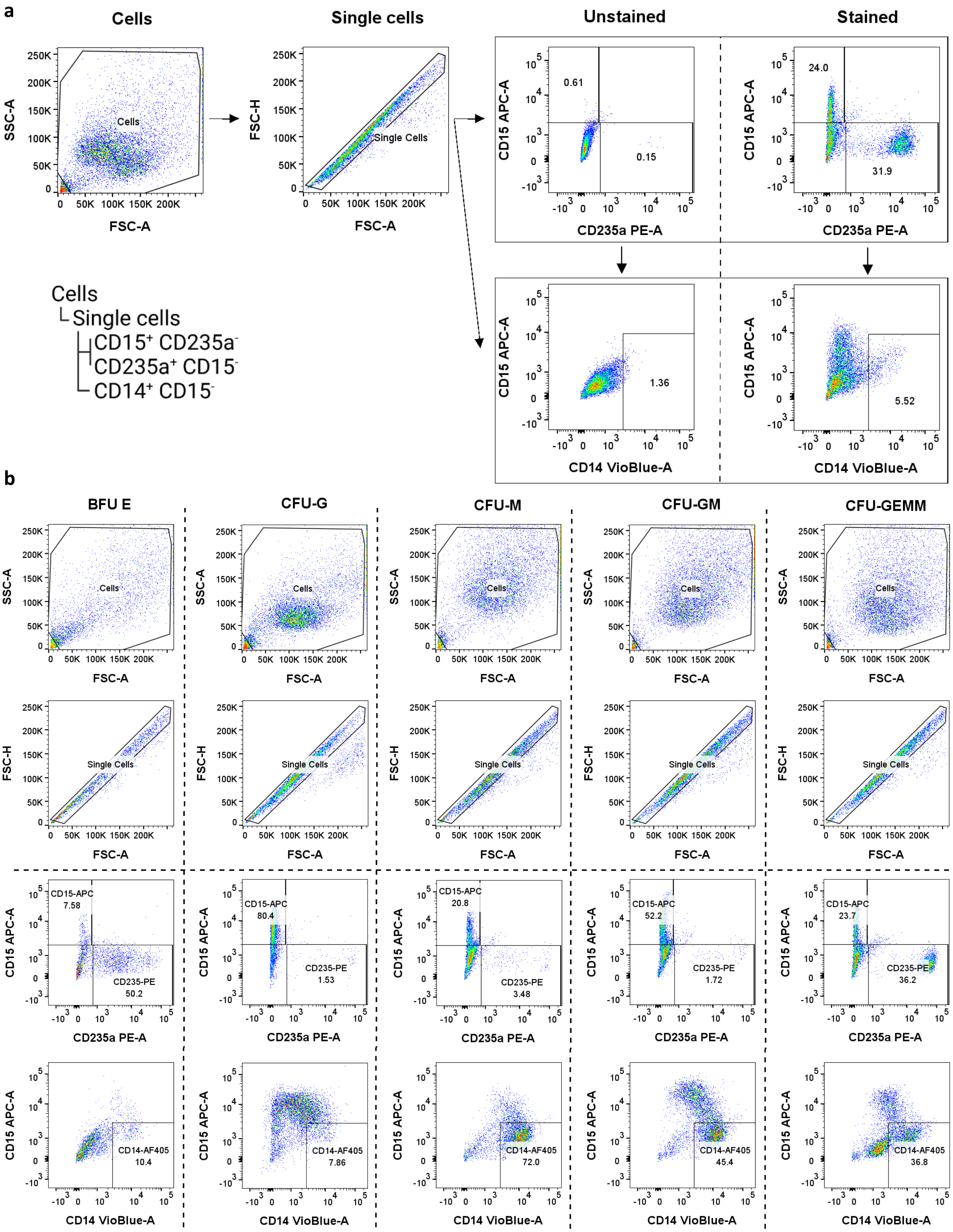


| Colony type | % of CD15+ cells | % of CD14+ cells | % ofCD235a cells (Glycophorin A cells) |
|---|---|---|---|
| CFU-GEMM | ≥15 | ≥15 | ≥20 |
| CFU-GM | ≥30 | ≥30 | |
| CFU-M | | ≥50 | |
| BFU-E | | | ≥50 |
| CFU-G | ≥50 | | |

a

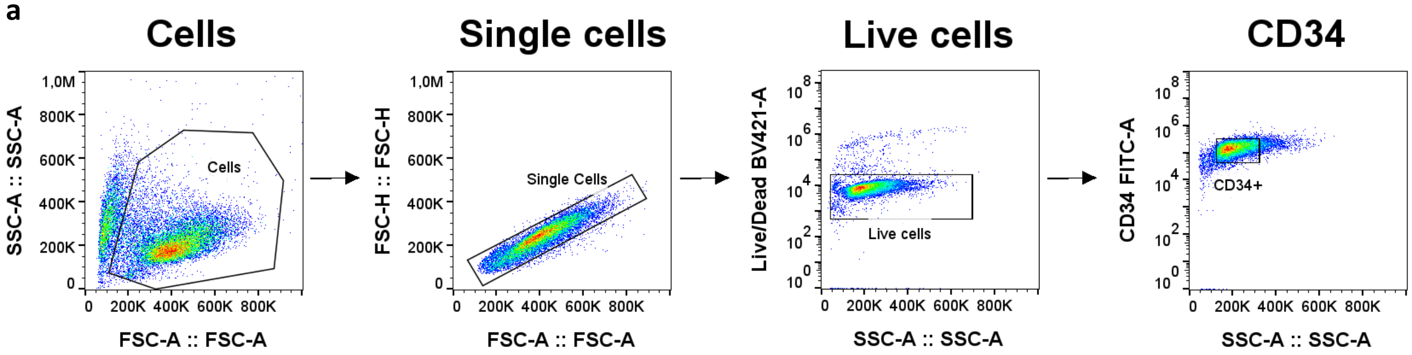
Cells
Single cells
Live cells
CD34
SSC-A :: SSC-A
FSC-A :: FSC-A
Cells
FSC-H :: FSC-H
Single Cells
Live/Dead BV421-A
Live cells
CD34 FITC-A
CD34+

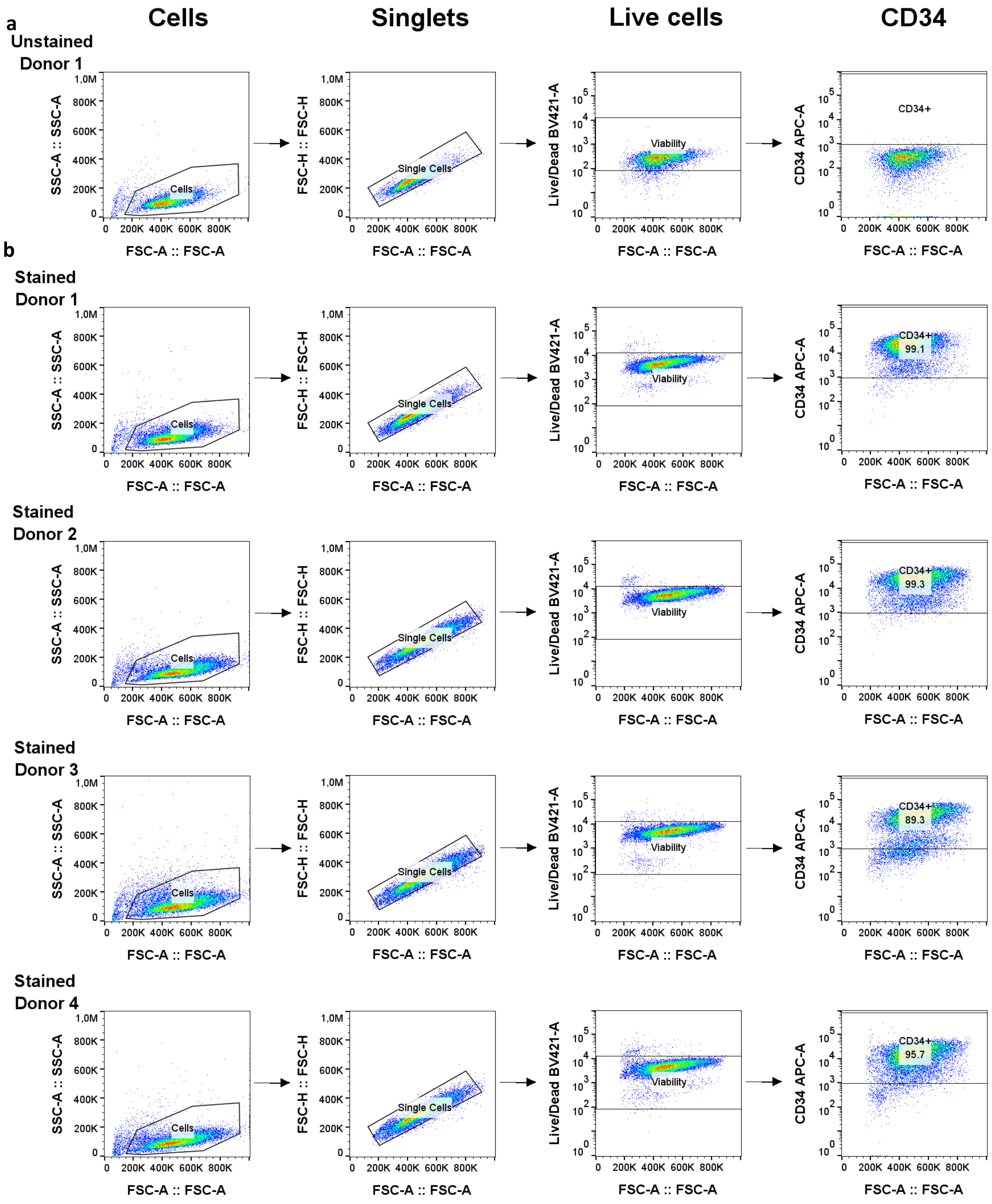

a
Cells
Singlets
Live cells
CD34
Unstained
Donor 1
b
Stained
Donor 1
Stained
Donor 2
Stained
Donor 3
Stained
Donor 4
SSC-A :: SSC-A
FSC-H :: FSC-H
Live/Dead BV421-A
CD34 APC-A
FSC-A :: FSC-A
Cells
Single Cells
Viability
CD34+
99.1
99.3
89.3
95.7

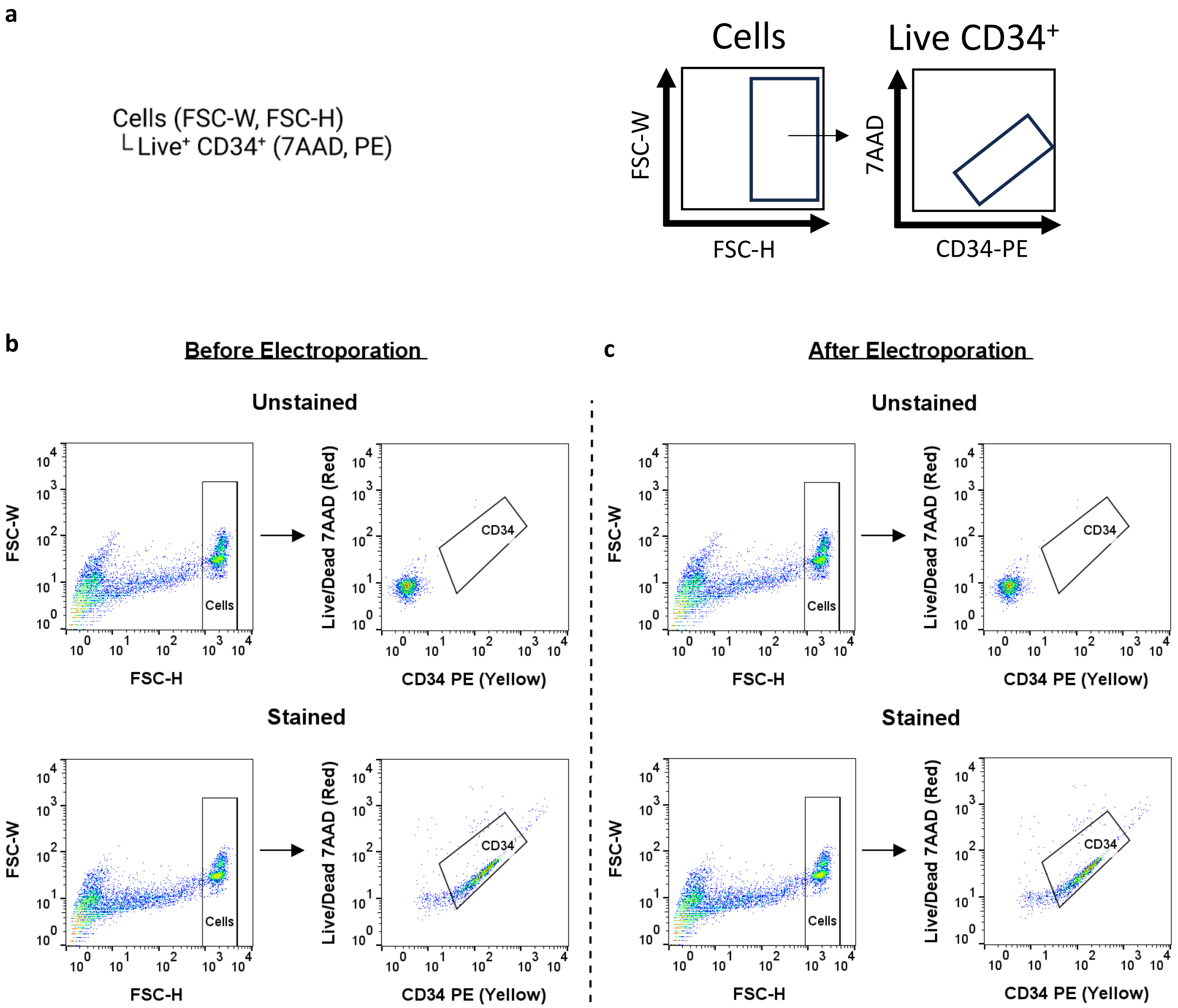
a
Cells (FSC-W, FSC-H)
└ Live⁺ CD34⁺ (7AAD, PE)
Cells
Live CD34⁺
FSC-W
FSC-H
7AAD
CD34-PE
b
Before Electroporation
c
After Electroporation
Unstained
Stained
FSC-W
FSC-H
Cells
Live/Dead 7AAD (Red)
CD34 PE (Yellow)
CD34

**a**

| Flow marker | Clone | Cat. # | Antibody type |
|---|---|---|---|
| PE anti-human CD33 | WM53 | 303404 | Mouse anti human |
| APC anti-human CD19 | HIB19 | 982406 | Mouse anti human |
| Alexa Fluor 700 anti-human HLA-A, B, C | W6/32 | 311438 | Mouse anti human |
| Pacific Blue anti-human CD45 | HI30 | 982306 | Mouse anti human |
| PE/Cyanine7 anti-mouse CD45.1 | A20 | 110730 | Mouse (A. SW) |
| PE/Cyanine5 anti-mouse TER-119 | TER-119 | 116210 | Rat anti mouse |
| FITC anti-human Lineage Coctail | RPA-2.10, OKT3, 61D3, CB16, HIB19, TULY56, HIR2 | 22-7778-72 | Mouse anti human |
| FITC-anti-human CD10 | HI10a | 312208 | Mouse anti human |
| APC anti-human CD34 | 561 | 343608 | Mouse anti human |
| PE/Cyanine7 anti-human CD38 | HIT2 | 303516 | Mouse anti human |
| Brilliant Violet 421 anti-human CD90 (Thy-1) | "5E10" | 328122 | Mouse anti human |
| PE/Cyanine5 anti-human CD45RA | HI100 | 304110 | Mouse anti human |
| Brilliant Violet 711 anti-mouse CD45.1 | A20 | 110739 | Mouse anti human |
| PE, anti-human CD45 | HI30 | 304039 | mouse anti human |

**b**

**Flow panel 1**

Cells
└ Single cells
  └ Live cells (Live/Dead Aqua)
    └ mTER11$^-$ (PE-Cy5)
      ├ mCD45.1$^+$ (mouse; PE-Cy7)
      └ hCD45$^+$ HLA A, B, C$^+$ (human; Pacific Blue, AF 700)
        ├ CD33$^+$ (myeloid cells; PE)
        └ CD19$^+$ (myeloid cells; APC)

**Flow panel 2**

Cells
└ Single cells
  └ Live cells (Live/Dead Aqua)
    ├ mCD45.1$^+$ (mouse; BV-711)
    └ hCD45$^+$ (human; PE)
      └ Lineage$^-$ (lineage exclusion; FITC)
        └ CD34$^+$ CD38$^-$ (HSPCs; APC; PE-Cy7)
          ├ CD90$^+$ CD45RA$^-$ (HSC; BV 421, PE-Cy5)
          ├ CD90$^-$ CD45RA$^-$ (MPP; BV 421, PE-Cy5)
          └ CD90$^-$ CD45RA$^+$ (LMPP; BV 421, PE-Cy5)

**a**

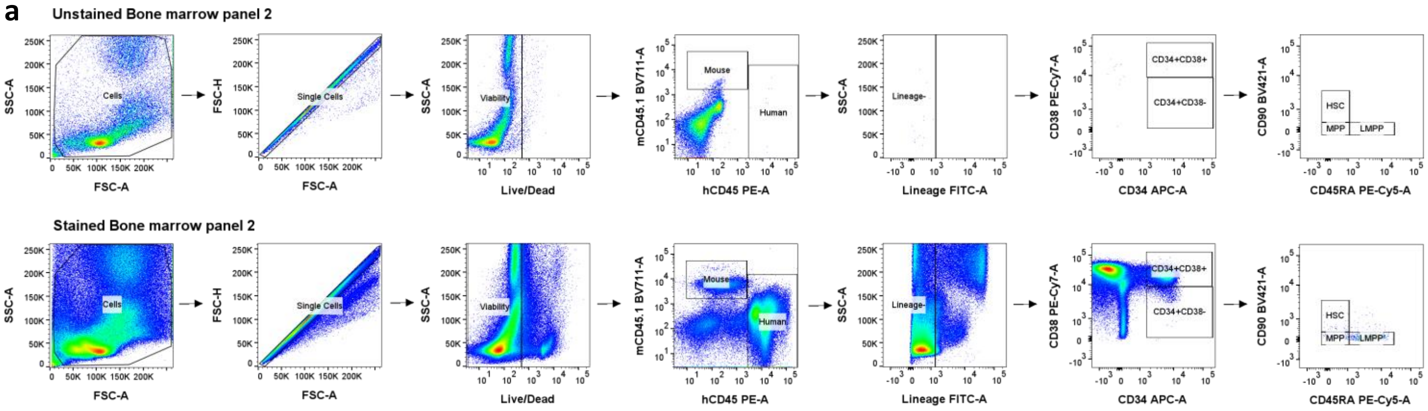
Unstained Bone marrow panel 2
Cells
Single Cells
Viability
Mouse
Human
Lineage-
CD34+CD38+
CD34+CD38-
HSC
MPP
LMPP
SSC-A
FSC-A
FSC-H
Live/Dead
mCD45.1 BV711-A
hCD45 PE-A
Lineage FITC-A
CD38 PE-Cy7-A
CD34 APC-A
CD90 BV421-A
CD45RA PE-Cy5-A
Stained Bone marrow panel 2

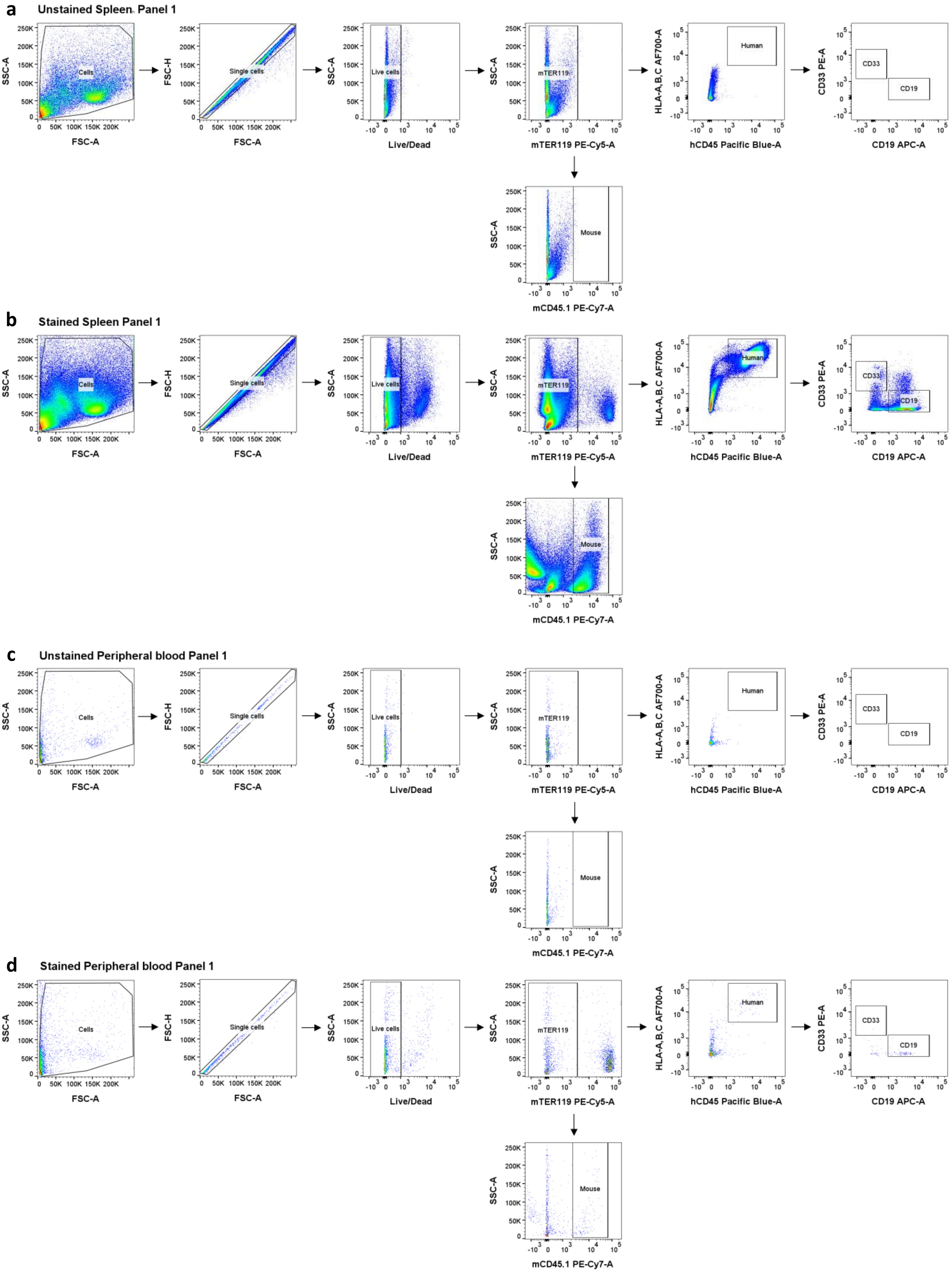
a
Unstained Spleen Panel 1
b
Stained Spleen Panel 1
c
Unstained Peripheral blood Panel 1
d
Stained Peripheral blood Panel 1
Cells
Single cells
Live cells
mTER119
Mouse
Human
CD33
CD19
SSC-A
FSC-A
FSC-H
Live/Dead
mTER119 PE-Cy5-A
mCD45.1 PE-Cy7-A
HLA-A,B,C AF700-A
hCD45 Pacific Blue-A
CD33 PE-A
CD19 APC-A

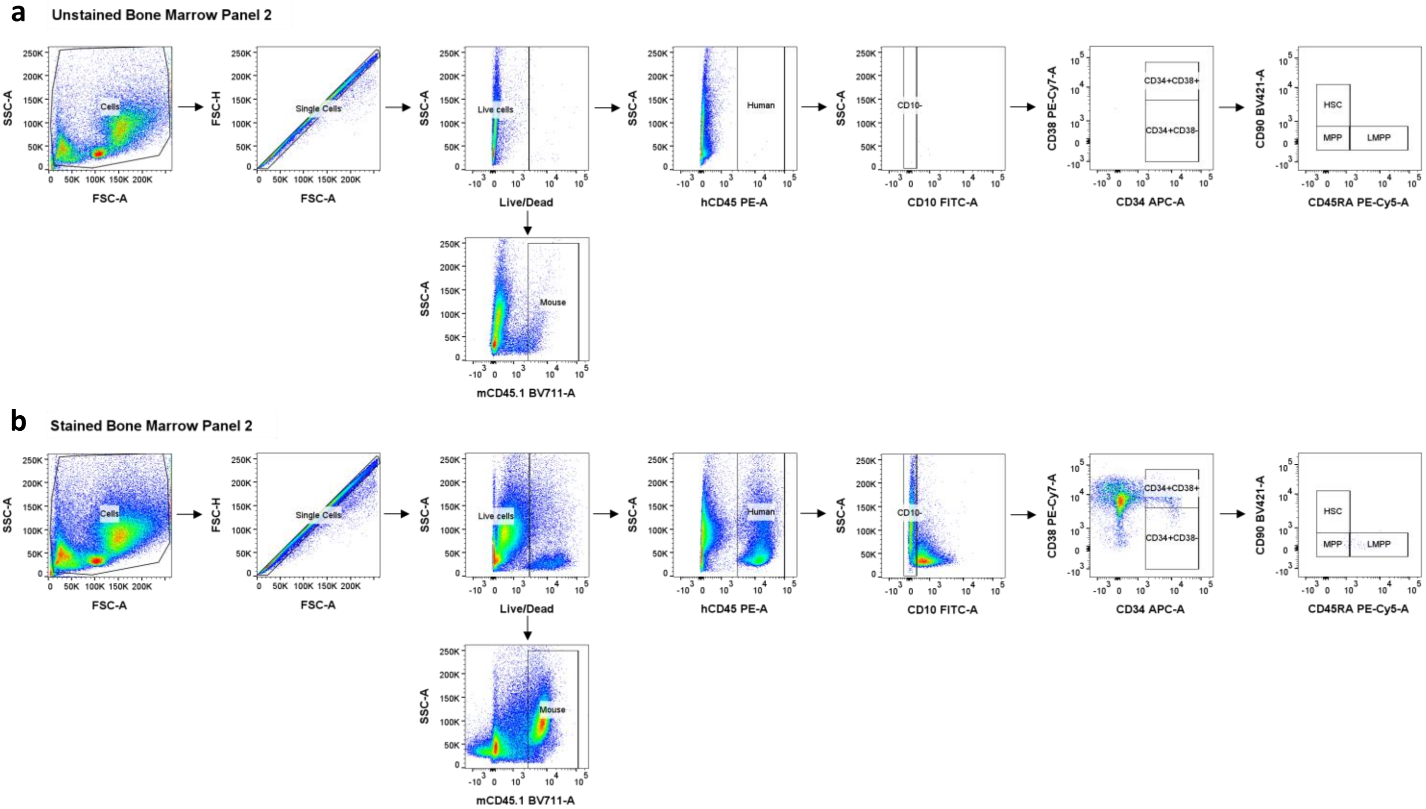
a
Unstained Bone Marrow Panel 2
Cells
Single Cells
Live cells
Human
CD10-
CD34+CD38+
CD34+CD38-
HSC
MPP
LMPP
Mouse
SSC-A
FSC-A
FSC-H
Live/Dead
hCD45 PE-A
CD10 FITC-A
CD38 PE-Cy7-A
CD34 APC-A
CD90 BV421-A
CD45RA PE-Cy5-A
mCD45.1 BV711-A
b
Stained Bone Marrow Panel 2
Cells
Single Cells
Live cells
Human
CD10-
CD34+CD38+
CD34+CD38-
HSC
MPP
LMPP
Mouse
SSC-A
FSC-A
FSC-H
Live/Dead
hCD45 PE-A
CD10 FITC-A
CD38 PE-Cy7-A
CD34 APC-A
CD90 BV421-A
CD45RA PE-Cy5-A
mCD45.1 BV711-A

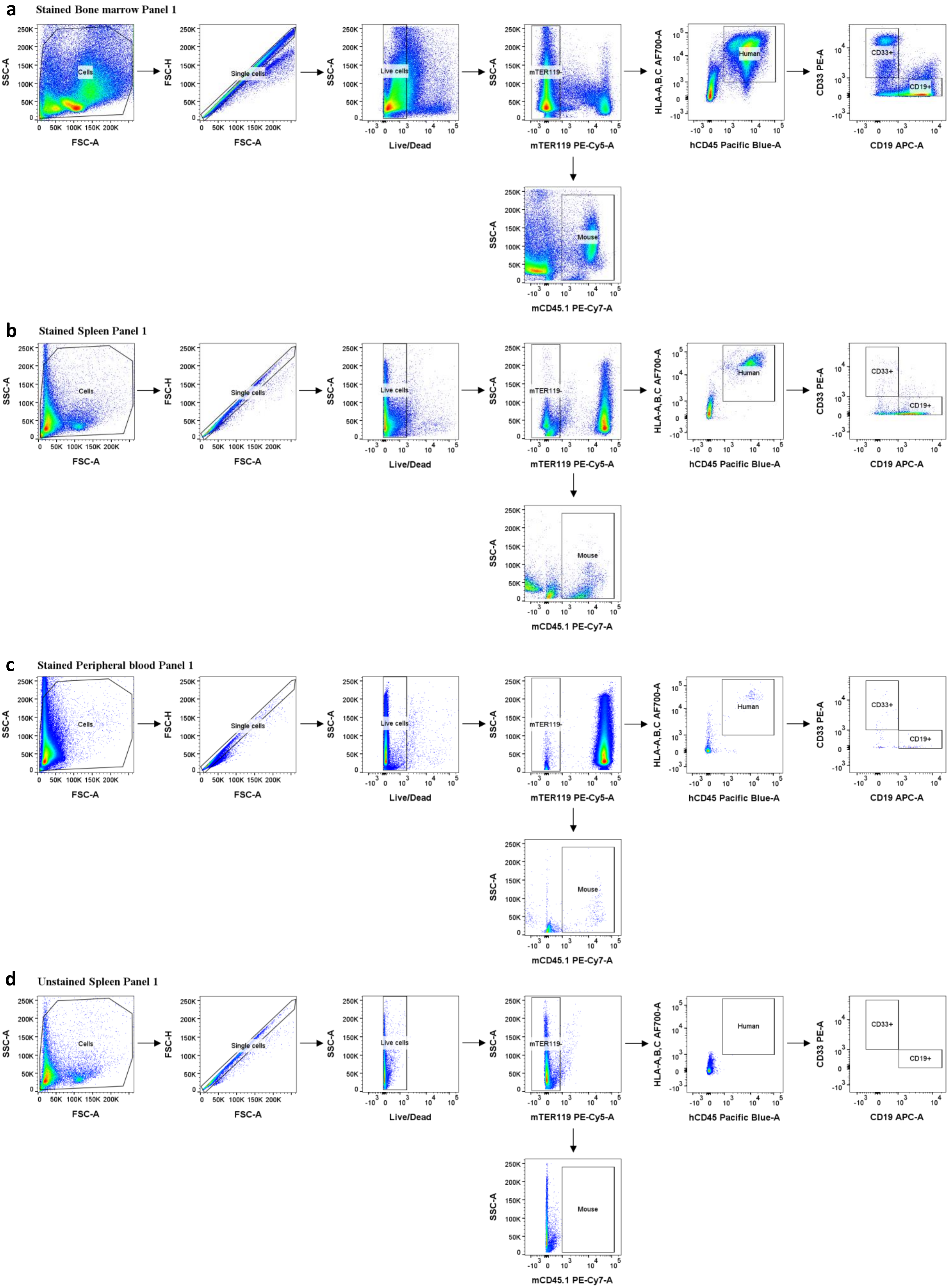
a
Stained Bone marrow Panel 1
b
Stained Spleen Panel 1
c
Stained Peripheral blood Panel 1
d
Unstained Spleen Panel 1
Cells
Single cells
Live cells
mTER119-
Human
CD33+
CD19+
Mouse
SSC-A
FSC-A
FSC-H
Live/Dead
mTER119 PE-Cy5-A
HLA-A,B,C AF700-A
hCD45 Pacific Blue-A
CD33 PE-A
CD19 APC-A
mCD45.1 PE-Cy7-A

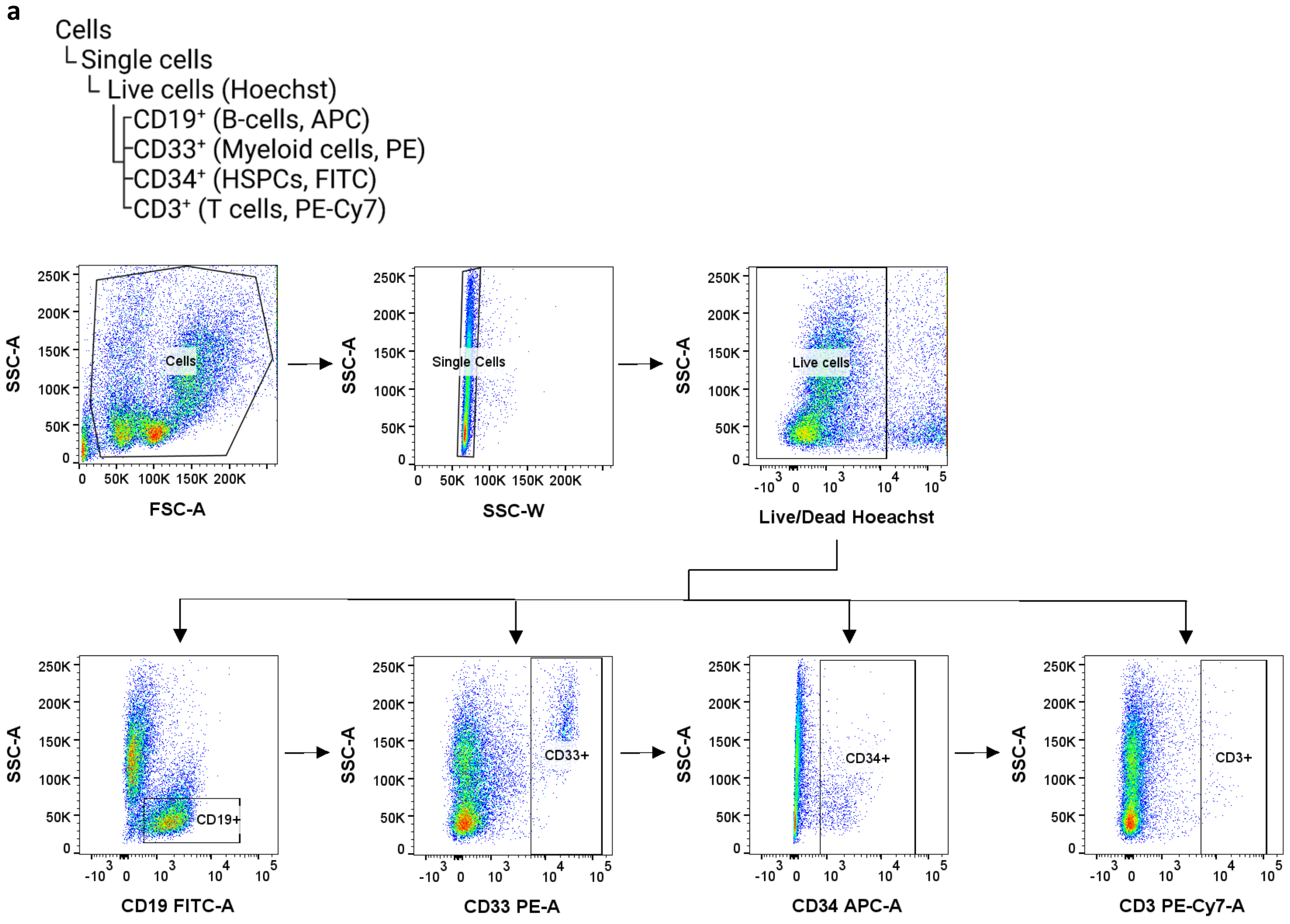
a
Cells
└Single cells
└ Live cells (Hoechst)
CD19+ (B-cells, APC)
CD33+ (Myeloid cells, PE)
CD34+ (HSPCs, FITC)
CD3+ (T cells, PE-Cy7)
Cells
Single Cells
Live cells
CD19+
CD33+
CD34+
CD3+
SSC-A
FSC-A
SSC-W
Live/Dead Hoeachst
CD19 FITC-A
CD33 PE-A
CD34 APC-A
CD3 PE-Cy7-A

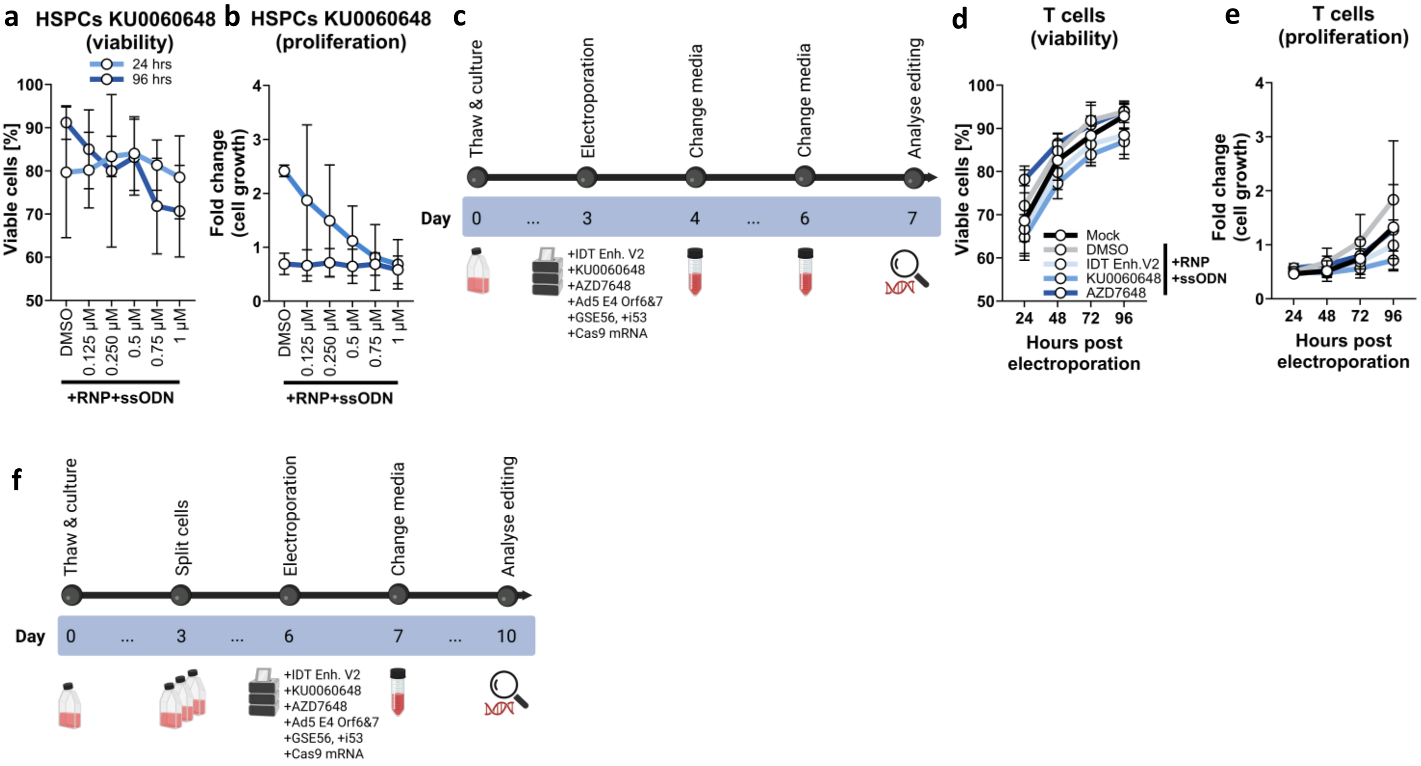

a
HSPCs KU0060648 (viability)
24 hrs
96 hrs
Viable cells [%]
100
90
80
70
60
50
DMSO
0.125 µM
0.250 µM
0.5 µM
0.75 µM
1 µM
+RNP+ssODN
b
HSPCs KU0060648 (proliferation)
Fold change (cell growth)
4
3
2
1
0
DMSO
0.125 µM
0.250 µM
0.5 µM
0.75 µM
1 µM
+RNP+ssODN
c
Thaw & culture
Electroporation
Change media
Change media
Analyse editing
Day
0
...
3
4
...
6
7
+IDT Enh. V2
+KU0060648
+AZD7648
+Ad5 E4 Orf6&7
+GSE56, +i53
+Cas9 mRNA
d
T cells (viability)
Viable cells [%]
100
90
80
70
60
50
24
48
72
96
Hours post electroporation
Mock
DMSO
IDT Enh.V2
KU0060648
AZD7648
+RNP +ssODN
e
T cells (proliferation)
Fold change (cell growth)
4
3
2
1
0
24
48
72
96
Hours post electroporation
f
Thaw & culture
Split cells
Electroporation
Change media
Analyse editing
Day
0
...
3
...
6
7
...
10
+IDT Enh. V2
+KU0060648
+AZD7648
+Ad5 E4 Orf6&7
+GSE56, +i53
+Cas9 mRNA

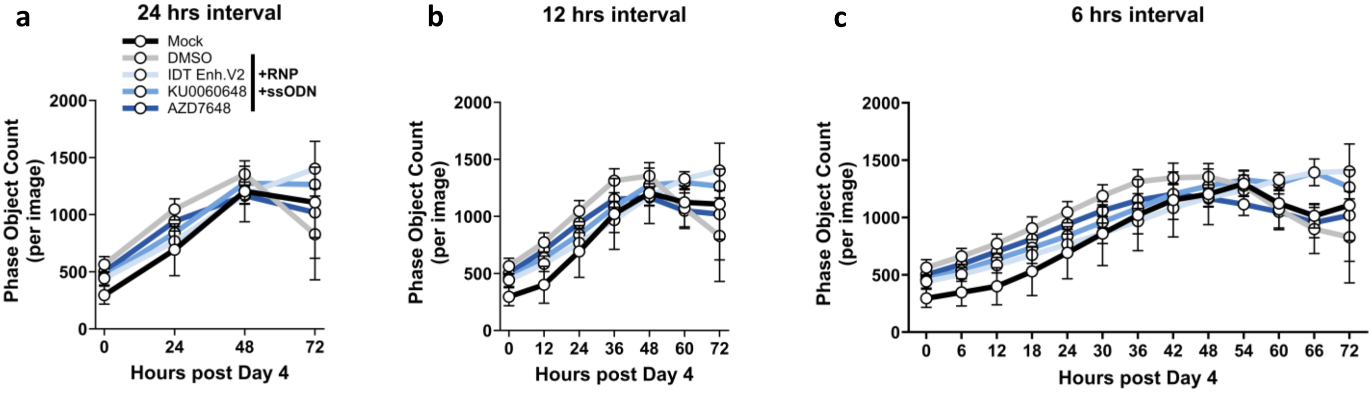

a
24 hrs interval
Mock
DMSO
IDT Enh.V2
KU0060648
AZD7648
+RNP
+ssODN
Phase Object Count (per image)
Hours post Day 4
b
12 hrs interval
Phase Object Count (per image)
Hours post Day 4
c
6 hrs interval
Phase Object Count (per image)
Hours post Day 4

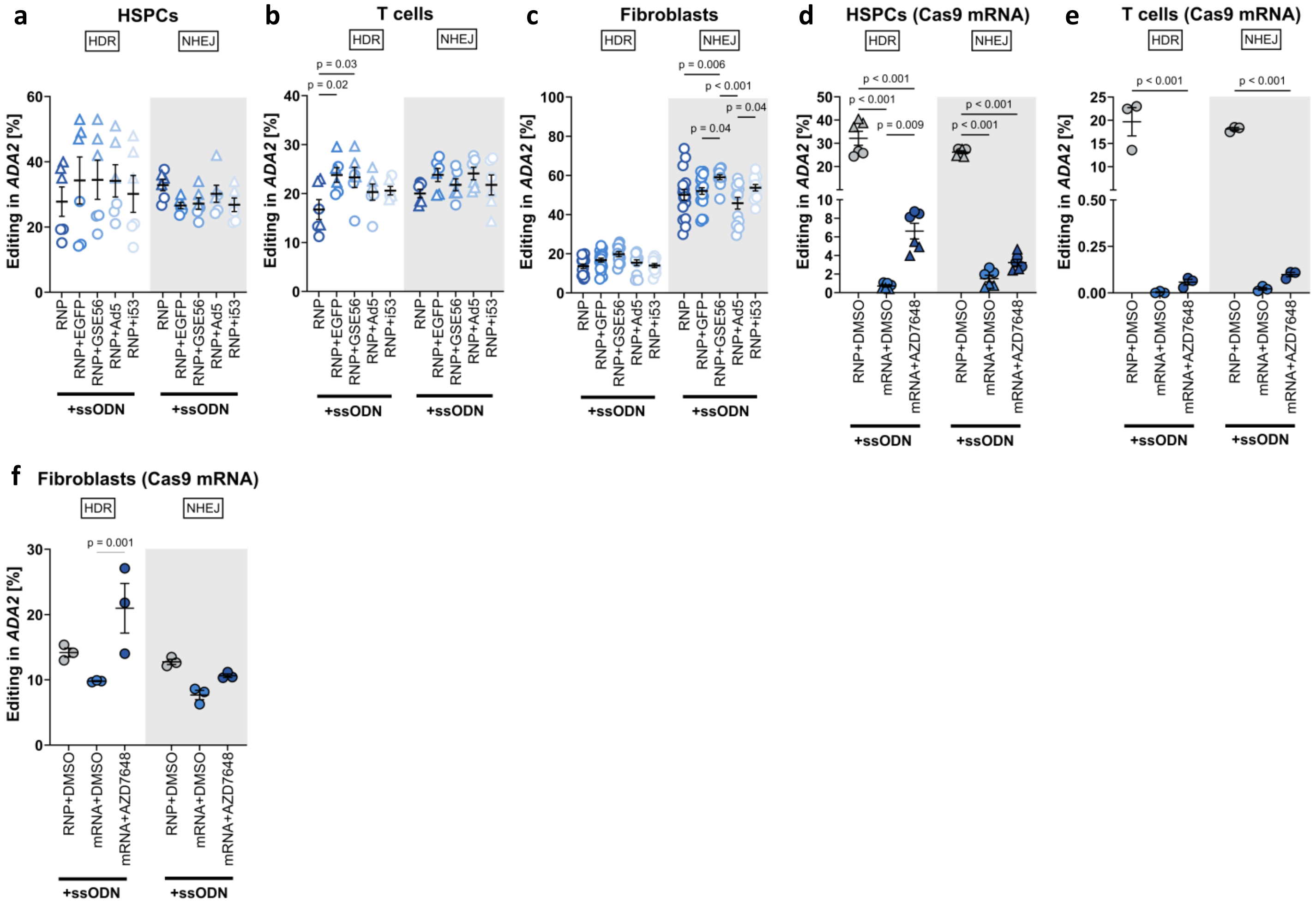
a
HSPCs
HDR
NHEJ
Editing in ADA2 [%]
RNP
RNP+EGFP
RNP+GSE56
RNP+Ad5
RNP+i53
+ssODN
b
T cells
HDR
NHEJ
p = 0.03
p = 0.02
Editing in ADA2 [%]
RNP
RNP+EGFP
RNP+GSE56
RNP+Ad5
RNP+i53
+ssODN
c
Fibroblasts
HDR
NHEJ
p = 0.006
p < 0.001
p = 0.04
p = 0.04
Editing in ADA2 [%]
RNP
RNP+GFP
RNP+GSE56
RNP+Ad5
RNP+i53
+ssODN
d
HSPCs (Cas9 mRNA)
HDR
NHEJ
p < 0.001
p < 0.001
p = 0.009
p < 0.001
p < 0.001
Editing in ADA2 [%]
RNP+DMSO
mRNA+DMSO
mRNA+AZD7648
+ssODN
e
T cells (Cas9 mRNA)
HDR
NHEJ
p < 0.001
p < 0.001
Editing in ADA2 [%]
RNP+DMSO
mRNA+DMSO
mRNA+AZD7648
+ssODN
f
Fibroblasts (Cas9 mRNA)
HDR
NHEJ
p = 0.001
Editing in ADA2 [%]
RNP+DMSO
mRNA+DMSO
mRNA+AZD7648
+ssODN

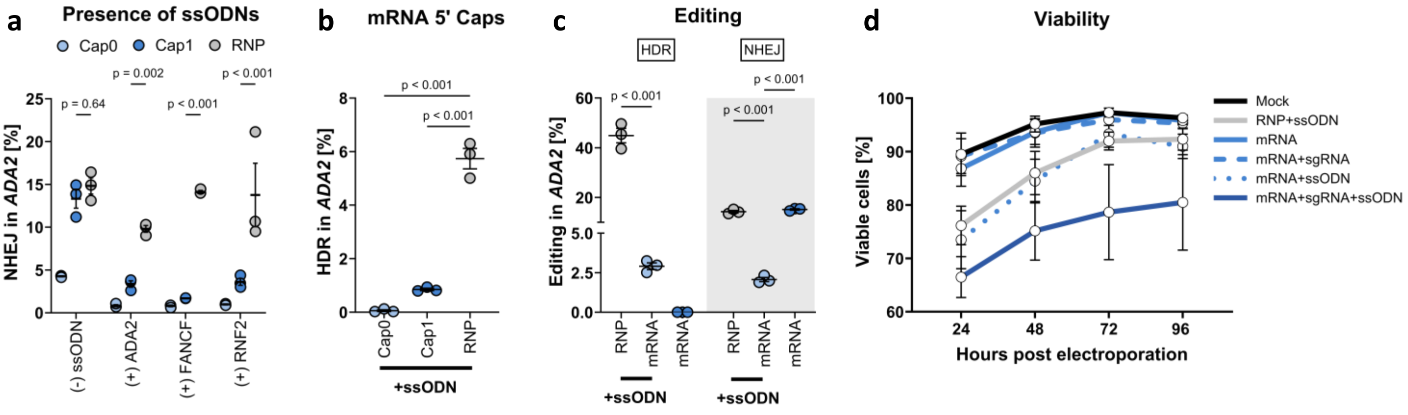
a
Presence of ssODNs
Cap0
Cap1
RNP
NHEJ in ADA2 [%]
p = 0.64
p = 0.002
p < 0.001
p < 0.001
(-) ssODN
(+) ADA2
(+) FANCF
(+) RNF2
b
mRNA 5' Caps
HDR in ADA2 [%]
p < 0.001
p < 0.001
Cap0
Cap1
RNP
+ssODN
c
Editing
HDR
NHEJ
Editing in ADA2 [%]
p < 0.001
p < 0.001
p < 0.001
RNP
mRNA
mRNA
RNP
mRNA
mRNA
+ssODN
+ssODN
d
Viability
Viable cells [%]
Mock
RNP+ssODN
mRNA
mRNA+sgRNA
mRNA+ssODN
mRNA+sgRNA+ssODN
Hours post electroporation

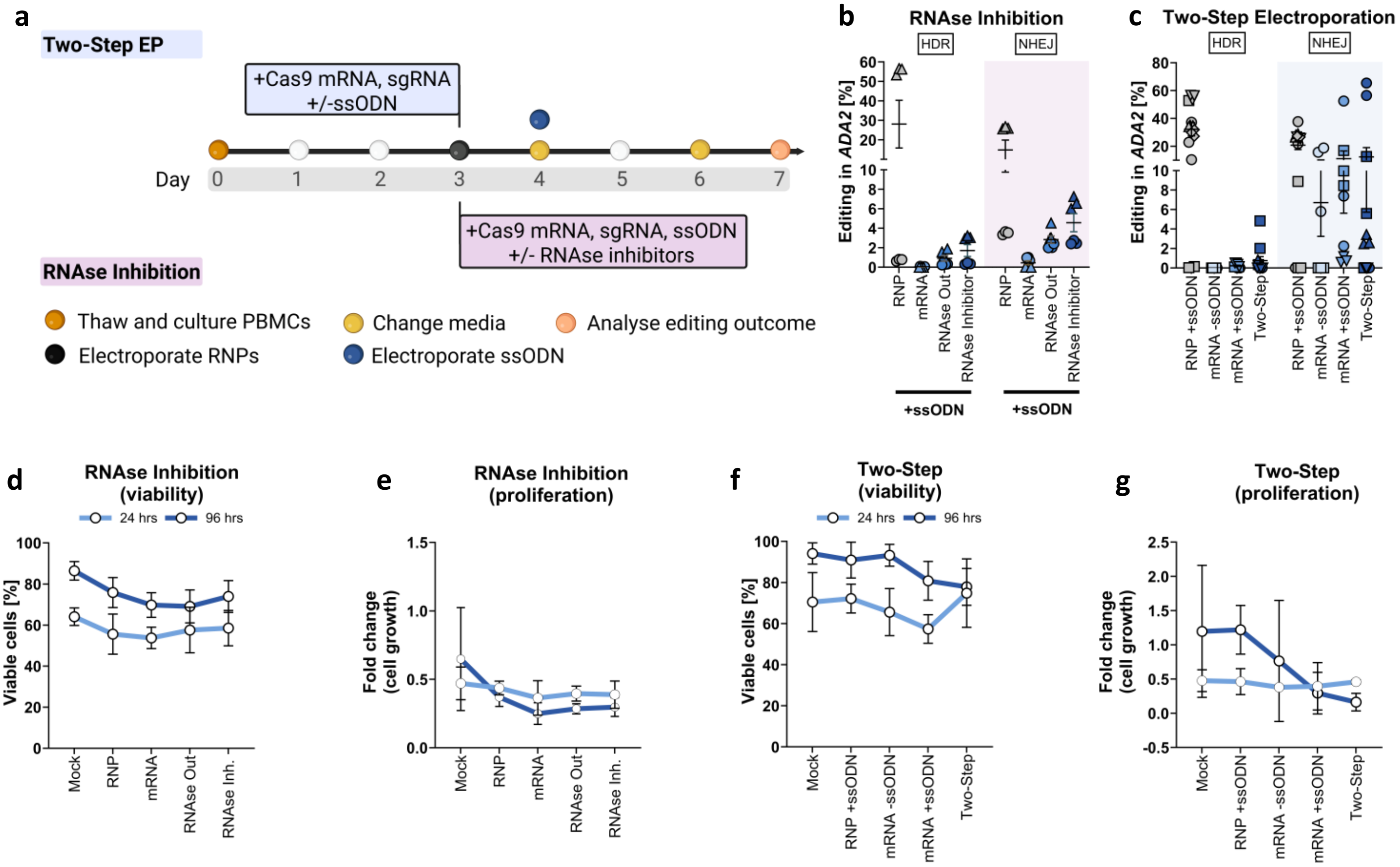
a
Two-Step EP
+Cas9 mRNA, sgRNA
+/-ssODN
Day
0
1
2
3
4
5
6
7
+Cas9 mRNA, sgRNA, ssODN
+/- RNAse inhibitors
RNAse Inhibition
Thaw and culture PBMCs
Electroporate RNPs
Change media
Electroporate ssODN
Analyse editing outcome
b
RNAse Inhibition
HDR
NHEJ
Editing in ADA2 [%]
RNP
mRNA
RNAse Out
RNAse Inhibitor
+ssODN
c
Two-Step Electroporation
RNP +ssODN
mRNA -ssODN
mRNA +ssODN
Two-Step
d
RNAse Inhibition
(viability)
24 hrs
96 hrs
Viable cells [%]
Mock
RNAse Inh.
e
RNAse Inhibition
(proliferation)
Fold change
(cell growth)
f
Two-Step
(viability)
g
Two-Step
(proliferation)

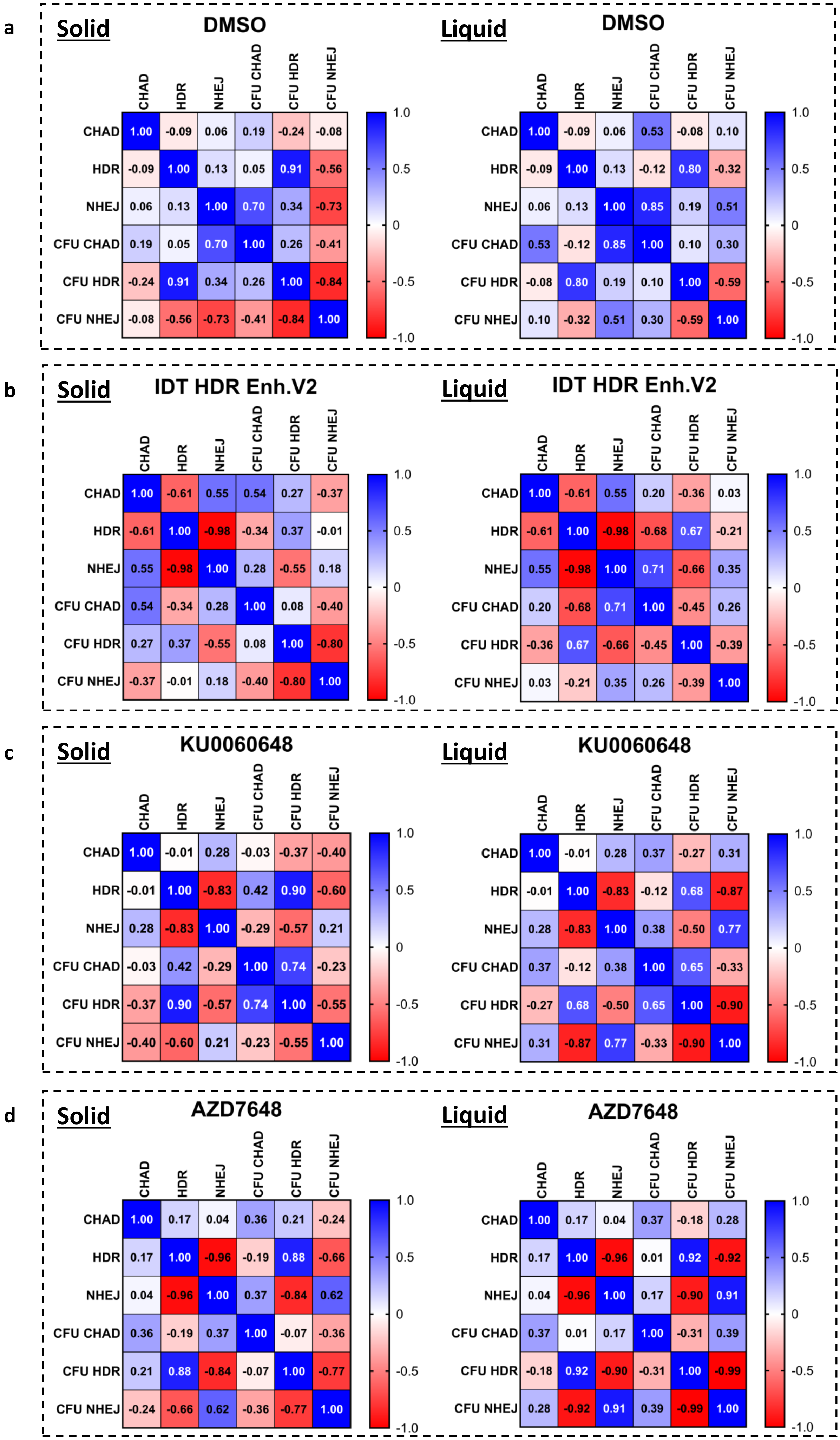
a
Solid
DMSO
CHAD HDR NHEJ CFU CHAD CFU HDR CFU NHEJ
CHAD 1.00 -0.09 0.06 0.19 -0.24 -0.08
HDR -0.09 1.00 0.13 0.05 0.91 -0.56
NHEJ 0.06 0.13 1.00 0.70 0.34 -0.73
CFU CHAD 0.19 0.05 0.70 1.00 0.26 -0.41
CFU HDR -0.24 0.91 0.34 0.26 1.00 -0.84
CFU NHEJ -0.08 -0.56 -0.73 -0.41 -0.84 1.00
1.0 0.5 0 -0.5 -1.0
Liquid
DMSO
CHAD HDR NHEJ CFU CHAD CFU HDR CFU NHEJ
CHAD 1.00 -0.09 0.06 0.53 -0.08 0.10
HDR -0.09 1.00 0.13 -0.12 0.80 -0.32
NHEJ 0.06 0.13 1.00 0.85 0.19 0.51
CFU CHAD 0.53 -0.12 0.85 1.00 0.10 0.30
CFU HDR -0.08 0.80 0.19 0.10 1.00 -0.59
CFU NHEJ 0.10 -0.32 0.51 0.30 -0.59 1.00
1.0 0.5 0 -0.5 -1.0
b
Solid
IDT HDR Enh.V2
CHAD HDR NHEJ CFU CHAD CFU HDR CFU NHEJ
CHAD 1.00 -0.61 0.55 0.54 0.27 -0.37
HDR -0.61 1.00 -0.98 -0.34 0.37 -0.01
NHEJ 0.55 -0.98 1.00 0.28 -0.55 0.18
CFU CHAD 0.54 -0.34 0.28 1.00 0.08 -0.40
CFU HDR 0.27 0.37 -0.55 0.08 1.00 -0.80
CFU NHEJ -0.37 -0.01 0.18 -0.40 -0.80 1.00
1.0 0.5 0 -0.5 -1.0
Liquid
IDT HDR Enh.V2
CHAD HDR NHEJ CFU CHAD CFU HDR CFU NHEJ
CHAD 1.00 -0.61 0.55 0.20 -0.36 0.03
HDR -0.61 1.00 -0.98 -0.68 0.67 -0.21
NHEJ 0.55 -0.98 1.00 0.71 -0.66 0.35
CFU CHAD 0.20 -0.68 0.71 1.00 -0.45 0.26
CFU HDR -0.36 0.67 -0.66 -0.45 1.00 -0.39
CFU NHEJ 0.03 -0.21 0.35 0.26 -0.39 1.00
1.0 0.5 0 -0.5 -1.0
c
Solid
KU0060648
CHAD HDR NHEJ CFU CHAD CFU HDR CFU NHEJ
CHAD 1.00 -0.01 0.28 -0.03 -0.37 -0.40
HDR -0.01 1.00 -0.83 0.42 0.90 -0.60
NHEJ 0.28 -0.83 1.00 -0.29 -0.57 0.21
CFU CHAD -0.03 0.42 -0.29 1.00 0.74 -0.23
CFU HDR -0.37 0.90 -0.57 0.74 1.00 -0.55
CFU NHEJ -0.40 -0.60 0.21 -0.23 -0.55 1.00
1.0 0.5 0 -0.5 -1.0
Liquid
KU0060648
CHAD HDR NHEJ CFU CHAD CFU HDR CFU NHEJ
CHAD 1.00 -0.01 0.28 0.37 -0.27 0.31
HDR -0.01 1.00 -0.83 -0.12 0.68 -0.87
NHEJ 0.28 -0.83 1.00 0.38 -0.50 0.77
CFU CHAD 0.37 -0.12 0.38 1.00 0.65 -0.33
CFU HDR -0.27 0.68 -0.50 0.65 1.00 -0.90
CFU NHEJ 0.31 -0.87 0.77 -0.33 -0.90 1.00
1.0 0.5 0 -0.5 -1.0
d
Solid
AZD7648
CHAD HDR NHEJ CFU CHAD CFU HDR CFU NHEJ
CHAD 1.00 0.17 0.04 0.36 0.21 -0.24
HDR 0.17 1.00 -0.96 -0.19 0.88 -0.66
NHEJ 0.04 -0.96 1.00 0.37 -0.84 0.62
CFU CHAD 0.36 -0.19 0.37 1.00 -0.07 -0.36
CFU HDR 0.21 0.88 -0.84 -0.07 1.00 -0.77
CFU NHEJ -0.24 -0.66 0.62 -0.36 -0.77 1.00
1.0 0.5 0 -0.5 -1.0
Liquid
AZD7648
CHAD HDR NHEJ CFU CHAD CFU HDR CFU NHEJ
CHAD 1.00 0.17 0.04 0.37 -0.18 0.28
HDR 0.17 1.00 -0.96 0.01 0.92 -0.92
NHEJ 0.04 -0.96 1.00 0.17 -0.90 0.91
CFU CHAD 0.37 0.01 0.17 1.00 -0.31 0.39
CFU HDR -0.18 0.92 -0.90 -0.31 1.00 -0.99
CFU NHEJ 0.28 -0.92 0.91 0.39 -0.99 1.00
1.0 0.5 0 -0.5 -1.0

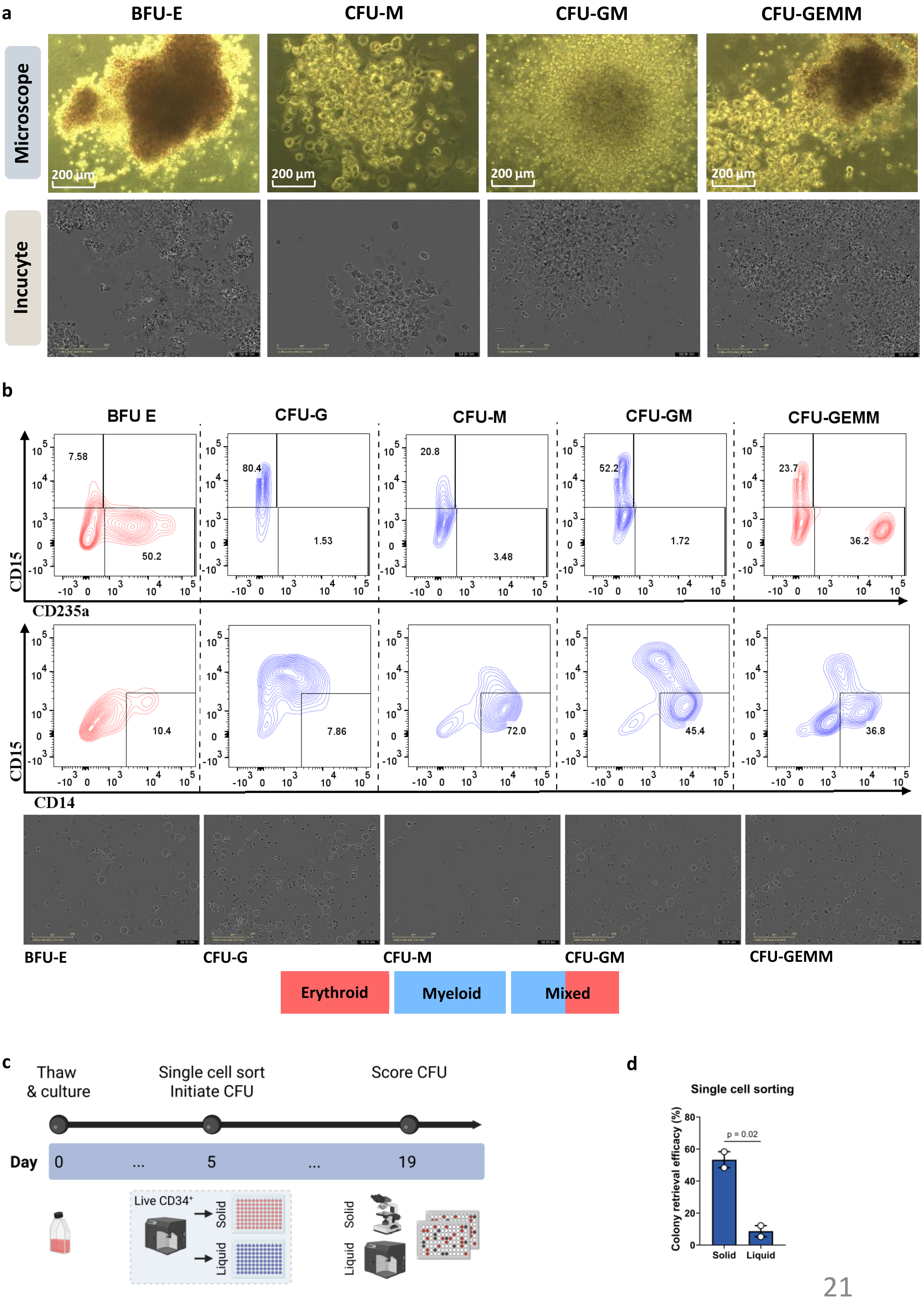
a
BFU-E
CFU-M
CFU-GM
CFU-GEMM
Microscope
Incucyte
200 µm
b
BFU E
CFU-G
CFU-M
CFU-GM
CFU-GEMM
7.58
50.2
80.4
1.53
20.8
3.48
52.2
1.72
23.7
36.2
CD15
CD235a
10.4
7.86
72.0
45.4
36.8
CD14
BFU-E
CFU-G
CFU-M
CFU-GM
CFU-GEMM
Erythroid
Myeloid
Mixed
c
Thaw
& culture
Single cell sort
Initiate CFU
Score CFU
Day
0
...
5
...
19
Live CD34+
Solid
Liquid
Solid
Liquid
d
Single cell sorting
Colony retrieval efficacy (%)
p = 0.02
Solid
Liquid

**a**

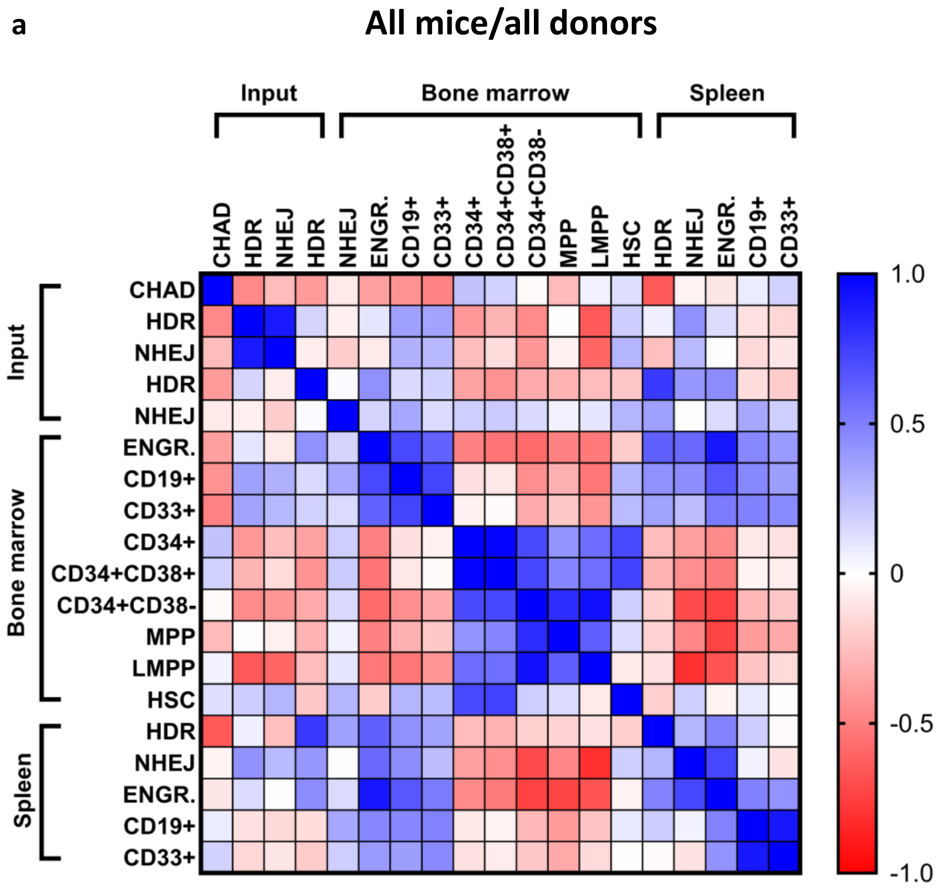
All mice/all donors
Input
Bone marrow
Spleen
CHAD
HDR
NHEJ
HDR
NHEJ
ENGR.
CD19+
CD33+
CD34+
CD34+CD38+
CD34+CD38-
MPP
LMPP
HSC
HDR
NHEJ
ENGR.
CD19+
CD33+
1.0
0.5
0
-0.5
-1.0

**b**

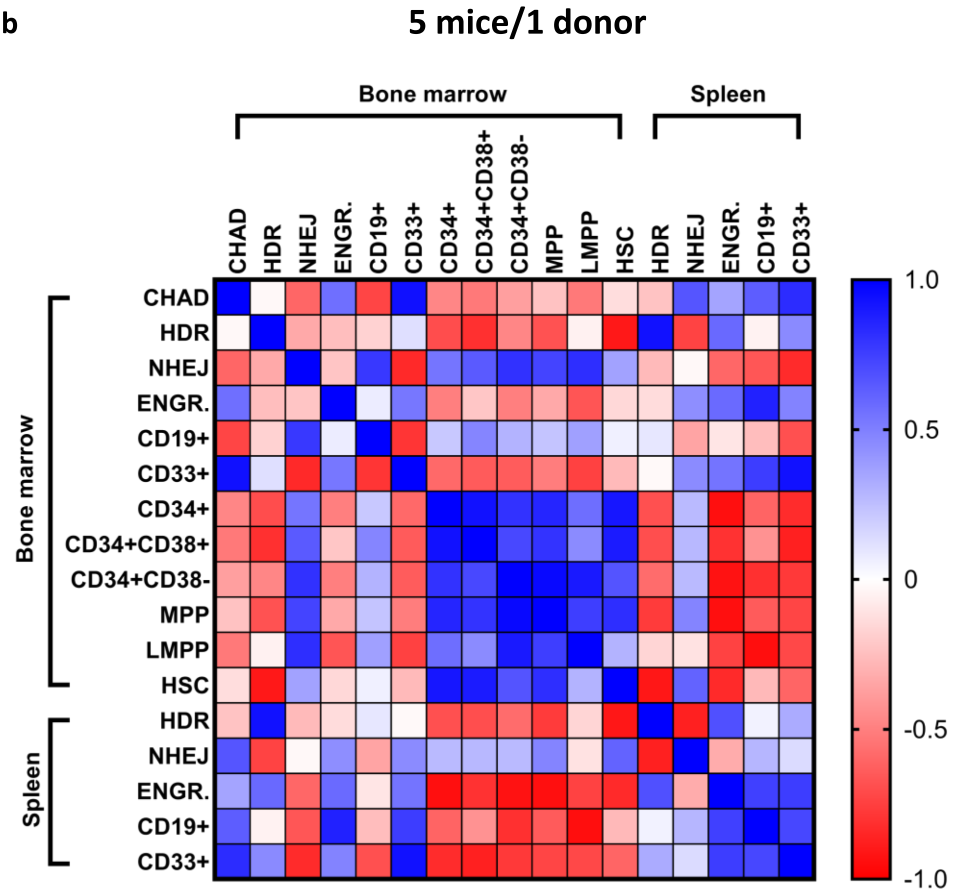
5 mice/1 donor
Bone marrow
Spleen
CHAD
HDR
NHEJ
ENGR.
CD19+
CD33+
CD34+
CD34+CD38+
CD34+CD38-
MPP
LMPP
HSC
HDR
NHEJ
ENGR.
CD19+
CD33+
1.0
0.5
0
-0.5
-1.0

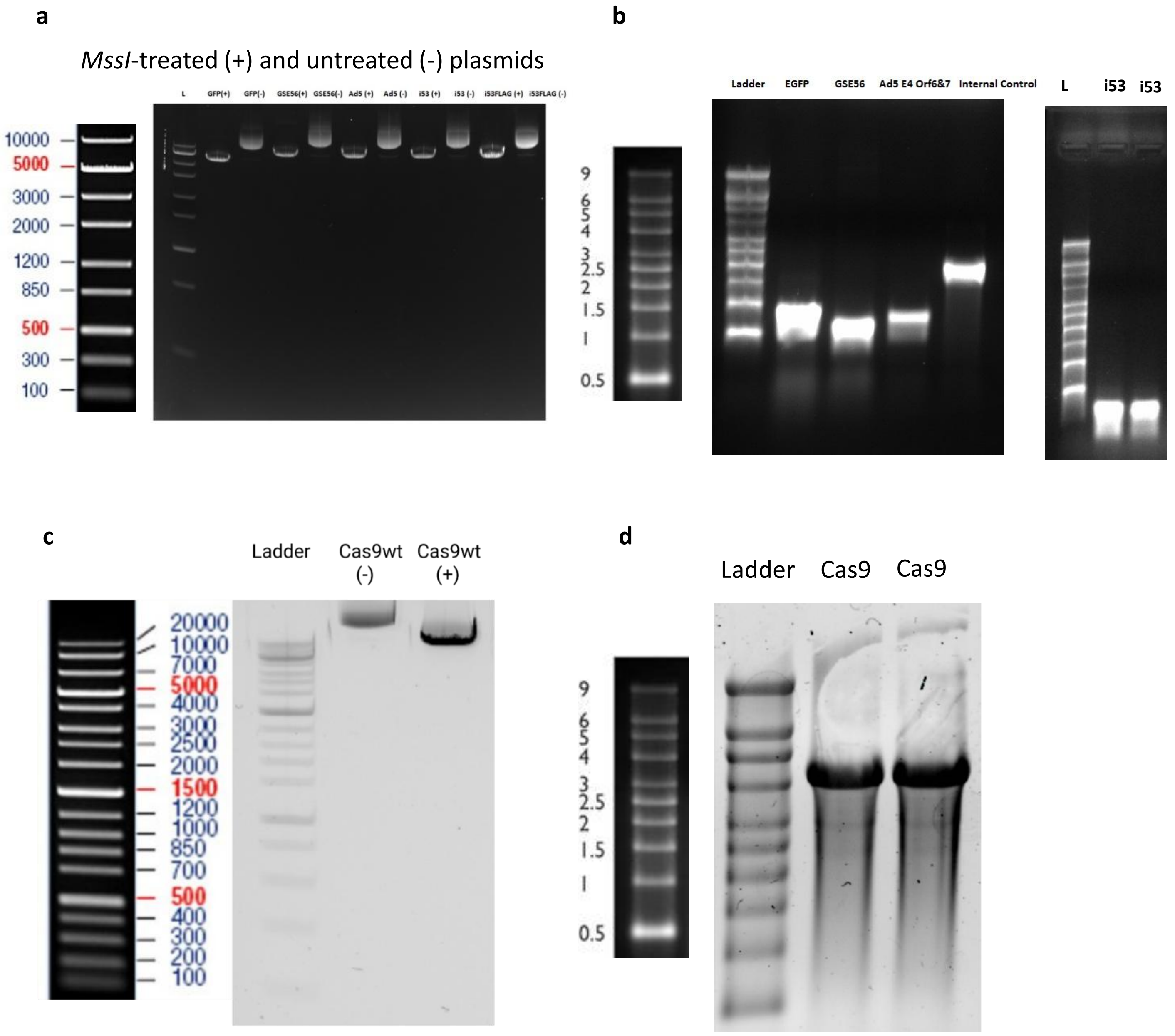
a
MssI-treated (+) and untreated (-) plasmids
L
GFP(+)
GFP(-)
GSE56(+)
GSE56(-)
Ad5 (+)
Ad5 (-)
i53 (+)
i53 (-)
i53FLAG (+)
i53FLAG (-)
10000
5000
3000
2000
1200
850
500
300
100
b
Ladder
EGFP
GSE56
Ad5 E4 Orf6&7
Internal Control
L
i53
i53
9
6
5
4
3
2.5
2
1.5
1
0.5
c
Ladder
Cas9wt (-)
Cas9wt (+)
20000
10000
7000
5000
4000
3000
2500
2000
1500
1200
1000
850
700
500
400
300
200
100
d
Ladder
Cas9
Cas9
9
6
5
4
3
2.5
2
1.5
1
0.5
