## Supplementary Material and Methods for "Editing of *ADA2* Point Mutation in Human Hematopoietic Stem Cells"

### Supplemental Methodology

#### Plasmid preparation

Coding sequences of *Cas9wt*[1], *GSE56*[2], *Ad5 Orf6&7*[3] and *i53*[4] were commercially synthesized and cloned into pENTR™ vector using GeneArt technology (Thermo Fisher). Next, fragments carrying the ORFs were subcloned into a T7-modifed pcDNA-DEST40™ (Thermo Fisher) backbone vector using Gateway system to obtain the expression vector. Briefly, each pENTR™ vector was mixed with pDEST40 in a 1:1 ratio, and 2 µl of LR Clonase enzyme (Thermo Fisher) was added, together with TRIS-EDTA (Invitrogen) to achieve a total reaction volume of 10 µl. The plasmid mix was incubated for 1 hour at room temperature before transformation into competent *E. Coli* dH5α (Thermo Fisher). pCMV-T7-EGFP was a gift from Benjamin Kleinstiver, Harvard University (Addgene plasmid #133962). The T7 promoter mutagenesis was performed by site-directed mutagenesis using Q5® Site-Directed Mutagenesis Kit (NEB) to enable the use of the CleanCap structure for *in vitro* transcription[5]. Briefly, a pair of primers was designed to introduce the desired nucleotide change, and the PCR reaction was carried out according to the manufacturer's instructions. After amplification, the samples were treated with KLD mix to ligate PCR products into circular form and degrade all unwanted byproducts. T7-modified mixture was then transformed into NEB 5-alpha Competent *E. Coli* provided with the kit. All plasmids were isolated and purified either by MiniPrep or Maxiprep kit (Qiagen) and were subjected to Sanger sequencing to verify correct cloning outcomes (Eurofins).

#### *In vitro* mRNA transcription

*In vitro* transcription (IVT) was carried out using either a T7 modified promoter for use with CleanCap or a T7 unmodified promoter to be used with Anti-Reverse Cap Analog (ARCA). 8 µg of each expression vector was linearised with 2 µl of Fast Digest *Mss*I restriction enzyme in FastDigest buffer (Invitrogen) in the final volume of 20 µl. Samples were incubated at 37°C overnight, followed by a heat inactivation at 65°C for 20 min. The linearisation was verified by gel electrophoresis using an E-gel (Invitrogen). For IVT with ARCA, 1 µg of linearised plasmid was mixed with 10 µl of 2xARCA/NTP mix (NEB) and 2 µl of T7 RNA Polymerase mix. The IVT reaction was incubated at 37°C for 30 min. Next, 2 µl of DNAse enzyme was added, and the mixture was incubated at 37°C for 15 min. The poly(A) tailing reaction was performed by adding 20 µl of RNAse-free water, 5 µl of 10X PolyA polymerase buffer, and 5 µl of 10X Poly(A) Polymerase (NEB) directly to the IVT reaction, followed by incubation at 37°C for 30 min. The final mRNA product was purified using LiCl solution, as described in the manufacturer's protocol. Aliquots were frozen in -80 °C until further use. For the IVT reaction with CleanCap, a similar protocol was followed with minor modifications. Firstly, UTP was replaced by N1-Methylpseudouridine (Trilink Biotechnologies). CleanCap was added according to the manufacturer's protocol. The IVT reaction was incubated at 37°C for 2 hrs followed by the same procedures as previously described for IVT reaction with ARCA. For all IVT reactions, RNAse-out (Thermo Fisher) was added to prevent mRNA degradation. The final mRNA products were evaluated on 1% denaturing agarose gel (with formaldehyde) and the final concentration was measured by Nanodrop (Fig E23 in the Online Repository).

**Genomic DNA isolation**

Genomic DNA isolation was performed using the DNA Blood & Tissue Kit (Qiagen). When QIAcube HT was employed, QIAAmp 96 DNA Kit was used, following the manufacturer's instructions. Briefly, cells were collected and centrifuged at 500 g for 5 min. The media was removed, and the cell pellet was washed in 500 µl of PBS and centrifuged at 500 g for 5 min. For the DNA elution step, 20-100 µl of elution buffer or MiliQ water was used. For the QIAcube, 90 µL of elution buffer was set as a default.

**Cell isolation**

Fibroblasts were obtained from skin biopsies, PBMCs were obtained from buffy coats and HSPCs were obtained from healthy donor cord blood.  For PBMCs and HSPCs enrichment, Lymphoprep and SepMate™-50 (IVD) (both STEMCELL Technologies) were used. For isolation of pure CD34+ population post enrichment, human CD34 MicroBead Kit UltraPure kit (Miltenyi Biotec) was utilised. Fibroblasts and PBMCs were cryopreserved in DMEM (Thermo Fisher) or RPMI 1640 (Thermo Fisher) supplemented with 10% DMSO. HSPCs were cryopreserved in CryoStor CS5 (STEMCELL Technologies). For long term storage, all cells were kept at -150 °C.

**Cell culture**

All cells were cultured at 37°C, with 21% $O_2$ and 5% $CO_2$ in a humidified incubator equipped with an open water reservoir. HSPCs were maintained in StemSpan™ SFEM II serum-free medium (STEMCELL Technologies), supplemented with recombinant human (Peprotech) thrombopoietin (TPO) (50 ng/ml), stem cell factor (SCF) (100 ng/ml), Flt3 ligand (Flt3-L) (100 ng/ml), and IL-6 (50 ng/ml). Subsequently, small molecules (STEMCELL Technologies) UM729 (50 µM) and SR-1 (10 µM) were introduced, along with 1% Pen-Strep (P/S) and GlutaMax™ (Gibco) (200mM) supplementation[3, 6]. Fibroblasts were maintained in DMEM (Thermo Fisher) supplemented with 10% FBS and with 1% P/S and were passaged upon reaching 80% confluency[7]. PBMCs were cultured in RPMI 1640 Medium (Thermo Fisher) with 10% FBS, supplemented with recombinant human (Peprotech) cytokines: IL-2 (120 U/ml), IL-7 (3 ng/ml), IL-15 (3 ng/ml), and ImmunoCult™ (STEMCELL Technologies) Human CD3/CD28 T Cell Activator (15 µl/ml)[7]. HSPCs were seeded at densities ranging from 100 000 - 500 000 cells/ml, for *in vivo* experiments, they were seeded at a density of 100 000 cells/ml. PBMCs, and fibroblasts were seeded at a density of 1 000 000 cells/ml. Cell count and viability were conducted using Trypan Blue exclusion (Thermo Fisher).

**Electroporation**

Electroporation was performed accordingly with previous studies[6, 7]. Briefly, either Cas9 protein (obtained from IDT or the protein facility at Karolinska Institute, Sweden) or mRNA (1 µg) were used in conjunction with sgRNA (IDT) or annealed gRNA (comprised of crRNA and tracrRNA; IDT). Chemically unmodified or 3'end chemically modified (2xPT (phosphothiorate) ssODN (Ultramer™ DNA Oligo, IDT) were also employed. RNP complexes were formed by mixing gRNA and Cas9 protein, which were then incubated for 15 min at 37°C. For electroporation of HSPCs, 100 000-500 000 cells were used per well, for T cells and fibroblasts, 1 000 000 cells were electroporated per well. 4D-Nucleofector® device (Lonza) was utilised. mRNA genetic enhancers (3 µg) were co-electroporated along with the editing components. Specific codes were used for each cell type: HSPCs – DZ-100, PBMCs – EO-115 and fibroblasts – CA-137. The P3 primary solution was selected for all cell types. P3 buffer or TheraPeak™ buffer (both from Lonza) were utilised to resuspend and electroporate cells. Immediately following electroporation, 80 µl of freshly prepared media, without P/S, was added to each well and incubated for 15 min in a humidified incubator. Prior to electroporation, cells were washed in PBS and subsequently placed into cell-specific media that did not contain P/S. For T cells, the media was supplemented only with 10% FBS and IL-2 (250 U/ml). The media for all cell types were replaced the following day with freshly prepared media.

**Two-Step electroporation and RNAse Inhibition**

The electroporation procedure was performed as described above, with minor modifications. For this two-step electroporation protocol, 2 000 000 PBMCs were used per well in a 12-well plate for T cells. Initially, the cells were electroporated with Cas9 mRNA and sgRNA, and 24 hrs after the first electroporation, the cells were subjected to a second electroporation using only the ssODN repair template. For experiments involving RNAse inhibitors, either 40 U/µl of RNAse-Out (Thermo Fisher) or 40 U/µl of Protector RNAse Inhibitor (Merck) was added to the cell mixture containing Cas9 mRNA, sgRNA, ssODN, and electroporation buffer, and gently mixed prior to transferring the cell mixture into the electroporation well. The post-electroporation treatment of the cells followed the same protocol as described above.

**NHEJ inhibitors and cells treatment**

KU0060648 (TargetMol) and AZD7648 (MedChemExpress) were resuspended in sterile DMSO according to the manufacturer's instructions when a ready-made version was not available (Alt® IDT HDR Enhancer V2, 690nM). Cells were cultured in the appropriate media specific to each cell type, without P/S, either supplemented with NHEJ inhibitor or in media containing an equivalent concentration of DMSO. At a minimum, 50 % of the original media was removed 24h post-electroporation, and freshly prepared media containing 1% P/S was added to the cell suspension.

**Droplet digital PCR for gene editing evaluation**

Droplet-digital PCR (ddPCR) was employed for the precise assessment of editing outcomes, as previously described[1, 7], with examples of gating strategies are illustrated (Fig E2 in the Online Repository). Briefly, DNA concentration was normalised to 1 – 8 ng/µl. The final volume of the ddPCR reaction was 20 µl and consisted of followed reagents for one reaction: 10 µl of 2xddPCR Supermix for Probes (Bio-Rad), 8 µl of normalised DNA, 0.18 µl of primer (IDT) forward and 0.18 µl primer reverse (each 900nM), 0.05 µl of reference probe (IDT) (FAM, 250nM) and 0.05 µl of NHEJ or HDR probe (HEX, 250 nM) followed by addition of 1 µl of MiliQ water to top the reaction volume. The NHEJ and HDR detection assays were conducted in two separate ddPCR reactions. Subsequently, each reaction was loaded into a sample well of an eight-well disposable cartridge (DG8™; Bio-Rad) together with 70 µl of droplet generation oil (Bio-Rad). ddPCR droplets were generated using a QX200™ Droplet Generator (Bio-Rad). The generated droplets were transferred into a 96-well PCR plate, heat-sealed with foil, and amplified using a conventional thermal cycler (Bio-Rad) under the following conditions: 1) 95 °C – 10 min, 2) 94 °C – 30 sec, 56 °C – 3 min, step repeated 42 times, 3) 98 °C – 10 min, 4) 4 °C – hold. The final reaction volume for ddPCR was 20 µl. The plate was read using a ddPCR reader (Bio-Rad), and data analysis was performed using QuantaSoft™ or QX Manager software (both from Bio-Rad). Both HDR and NHEJ will lead to drop-off events. NHEJ is calculated by subtracting HDR values from total editing values. The formulas to calculate are as follows:

$$\%\,HDR = \left(\frac{Ref(+)HDR(+)}{Ref(+)\ HDR(+) + Ref(+)\ HDR(-)}\right) * 100$$

$$\%\,Total\ editing = \left(\frac{Ref(+)NHEJ(-)}{Ref(+)\ NHEJ(+) + Ref(+)\ NHEJ(-)}\right) * 100$$

$$\%\,NHEJ = Total\ editing\ (\%) - HDR\ (\%)$$

**ddPCR chromosomal arm deletion (CHAD) assay**

Chromosomal loss assays were designed to detect long deletions and losses of long arm distal from 22q11.1 locus, where *ADA2* is located. Loss probes (FAM) targeting deletions and amputations were developed to target sequences either 100bp or 500 kbp distally upstream of the CRISPR/Cas9 cutting side in *ADA2* gene towards the telomere. A reference probe was designed for *CASC11* gene, located on chromosome 8q24.21. Examples of gating strategies are presented (Fig E3, 4 in the Online Repository). DNA concentration was normalised to 8 ng/µl. The final volume of ddPCR reaction was 20 µl, which consisted of the followed reagents for a single reaction: 10 µl of 2xddPCR Supermix for Probes (Bio-Rad), 8 µl of normalised DNA, forward and reverse primers for reference and loss (each 450nM), a reference probe (HEX, 250nM) and a deletion or amputation probe (FAM, 250 nM). Additionally, 25 U of *Hind*III restriction enzyme (NEB) was incorporated into the master mix to digest the DNA into smaller fragments during oil droplet generation. MiliQ water was utilised to achieve the required final reaction volume. Deletion and amputation assays were conducted in parallel in different plates. The droplet generation, PCR amplification and ddPCR readout were perform as described above. Comparison requires normalising the CHAD level to unedited cells from the same donor, both *in vitro* and *in vivo* comparison. To calculate the percentage of chromosomal arm deletions, the formula provided below was employed[8]:

$$\%\ of\ loss = 100 * (1 - \frac{Target\ FAM(+)}{Reference\ HEX(+)})$$

**Colony forming unit (CFU) assay**

Semi-solid methylcellulose (STEMCELL Technologies) and liquid (Miltenyi Biotec) colony-forming unit (CFU) assays were performed according to manufacturer's instructions. For semi-solid CFU assay, 500 - 3000 cells were seeded to semi-solid media (1.1 ml) in duplicates into 35 mm dishes. The readout was performed by assessment of colony types based on microscopic features. For the liquid CFU assay, cells were diluted to 0.3 cells per well and distributed in 60 µl into 96-well plate using a multichannel pipette, with three replicates (3x96-well plates). The plates were placed in humidified chambers containing open 35 mm dishes filled with PBS to prevent evaporation. In addition, 500 cells were seeded into liquid media in a 12-well plate (1 ml). This culture was utilised for setting gates for subsequent flow cytometry analysis and genomic DNA isolation. Liquid CFU plates were assessed by flow cytometry 14 days after initiation according to manufacturer's protocol using a premixed antibody cocktail containing CD235a-PE, CD14-VioBlue® and CD15-APC (Miltenyi Biotec). Secondary solid CFU assay was performed as described elsewhere[9]. Briefly, cells were collected from 35 mm dishes by resuspending the cells embedded in the methylcellulose matrix in pre-warmed PBS. The cells were centrifuged at 300 g for 5 min and resuspended in 5 ml of PBS. A total of 5 000 cells were seeded (10x more than for the 1st CFU) into freshly thawed semi-solid CFU media (1.1 ml) and distributed into 35 mm dish in duplicates. For low-density liquid CFU culture, cells were passaged from the original culture at day 14 and reseeded at a density of 5 000 cells per 1 ml of liquid media in 12-well plate. For the single-cell sorting procedure, cells were stained for live $CD34^+$ in 100 µl volume of PBS (1 µl Live/Dead Violet (Thermo Fisher), 5 µl of CD34-FITC (BioLegend)) and blocked with 5 µl Human TruStain FcX™ Blocker (BioLegend). Sorted cells were deposited into 96-well plates (Fisher Scientific) containing 100 µl of semi-solid methylcellulose CFU media or 60 µl of liquid CFU media, utilising the SONY SH800 sorter. All related gating plots are shown (Fig E5, 6 and Table 1 in the Online Repository).

**CD34+ staining**

To detect CD34 on the surface of HSPCs treated with NHEJ inhibitors, the cells were stained with 2 µl of 7AAD (BioLegend) and 1 µl of CD34-PE (BioLegend) in a total volume of 100 µl of PBS and incubated in the dark at room temperature for 20 min. The cells were blocked at the same time with 5 µl of Human TruStain FcX$^{TM}$ Blocker. Samples were analysed using the Muse system (Cytek Biosciences) according to the manufacturer's protocol. Additionally, for the analysis of the CD34 marker, flow cytometry was performed utilising 1 µl Live/Dead Violet (Thermo Fisher) for viability staining and 5 µl of CD34-APC (BioLegend) in 100 µl of PBS. Cells were consistently blocked with 5µl of Human TruStain FcX$^{TM}$ Blocker (BioLegend). Samples were processed on the Sony SH800. Both Muse and flow cytometry data were analysed using FlowJo v10.10.0 software. The gating strategies are illustrated (Fig E7, 8 in the Online Repository).

**Incycute imaging**

HSPCs and CFU-derived cells were imagined by Incucyte system in both adherent and non-adherent modes. The analysis was conducted with Incucyte® Cell-by-Cell Analysis Software Module (Sartorius). Briefly, the cells were placed in an imaging closed culture unit and maintained at 37°C, with 21% $O_2$ and 5% $CO_2$ in a humidified incubator containing an open water reservoir. Individual image frames were selected to represent entire wells and compiled in the video files or as an individual photograph deposited online (https://osf.io/fkdh6/?view_only=99b31cdffbbe467b917bd60af2f613ae).

**Mice housing**

All animals were housed in the individual ventilated cages (IVC) provided with environmental enrichment (beddings, cottons, chewing wooden sticks, plastic houses and tubes). Mice had access to unlimited chow food supplemented with sunflower seeds, recovery gels, and acidified water (HCl based). The animals were maintained under a 12:12 light/dark cycle with controlled temperature conditions. Protocols for the humane euthanasia of experimentally ill animals were strictly adhered to, along with daily monitoring. Deceased mice that occurred during the experimental period prior to reaching the humane endpoint were excluded from data analysis.

#### Humanised mice

8 - 12 weeks old NOD.Cg-*Prkdc*$^{scid}$ *Il2rg*$^{tm1Wjl}$/SzJ (Charles River) male and female mice were sub-lethally irradiated with two doses of 1.25 Gy, separated by four hrs between each dose. For irradiation, CIX3 320kV Self-Contained Cabinet Irradiator with 300kV x 10,0mA was used. For 1.25Gy dose, the mice were exposed to X-rays for 1 min 34 sec utilising 700mm FSD level allowing two individually ventilated cages (IVC) being irradiated at the same time. Within 24 hrs post the $2^{nd}$ dose of irradiation, 200 000 human HSPCs, resuspended in a total volume of 200 µl of sterile PBS, were injected intravenously via tail vein into pre-warmed mice. Control mice were injected with a sterile 200 µl PBS solution. Mice were sacrificed 12- or 16-weeks post-irradiation by cervical dislocation. Mice injected only with PBS without irradiation served as controls for environment cleanliness and injections factor. Sentinel mice were housed in the same room. Organs were collected and stained for flow cytometry according to previously established protocols[6, 10]. All antibodies employed in this study are listed (Fig E9 and Table 2 in the Online Repository), while the gating plots are also presented (Fig E10-14 in the Online Repository). The cells were blocked with 5 µl of Human TruStain FcX$^{TM}$ Blocker (BioLegend) 5 µl of True-Stain Monocyte Blocker$^{TM}$ (BioLegend) in total volume of 100 µl. Amount of antibodies and Live/Dead Aqua viability dye for staining were used according to the manufacturer's protocols. Samples were analysed on BD® LSR II Flow Cytometer (BD Bioscience). For cell sorting, samples were blocked with 5 µl Human TruStain FcX$^{TM}$ Blocker (BioLegend) and stained with 5 µl CD19-FITC, 5 µl CD33-PE, 5 µl CD34-APC and 5 µl CD3-PE-Cy7 (all BioLegend) in total volume of 100 µl. Hoechst (1 µl) (Thermo Fisher) was used to stain for viable cells. Cell sorting was performed using BD® FACS Aria II Sorter (BD Biosciences). For flow cytometry data analysis, Kaluza 2.1 or FlowJo v10.10.0 software was employed.

#### Statistics

Data were analysed using unpaired t-test, One-way or Two-way ANOVA tests, followed by multiple comparison tests (Tukey's, Dunnett's or Šídák's). For *in vitro* experiments, a p-value <0.05 was considered statistically significant, while for *in vivo* experiments, a p-value of <0.10

was deemed significant due to the smaller number of animals used. To account for variability in cell viability and proliferation data, standard deviation (±SD) was employed, whereas for the remaining data comparing mean values, standard error of the mean (±SEM) was used. Tables containing all p-values are included in the corresponding statistic Excel files.
